# Osteoprotegerin-Enabled Immune Evasion of Pathological Adipose Stromal Cells Drives Metabolic Dysfunction in Obesity

**DOI:** 10.64898/2026.01.12.698936

**Authors:** Hara Apostolopoulou, Yao Wang, W. Reid Bolus, Radha Singh, Arjan Bains, Priyadarshini Bahadure, Kavya Gupta, Suneil K. Koliwad, Anil Bhushan

**Affiliations:** Diabetes Center, University of San Francisco, San Francisco CA, 94143, USA; Department of Medicine, University of San Francisco, San Francisco CA, 94143, USA

**Author notes:** Correspondence: Suneil K. Koliwad, M.D., Ph.D., University of California San Francisco, S1230A Medical Sciences, 513 Parnassus Ave., San Francisco, CA 94143-0669, Anil Bhushan, Ph.D., University of California San Francisco, 1045 Health Sciences West, San Francisco, CA 94143-0669. equal contribution (Suneil K. Koliwad, Anil Bhushan).

## Abstract

Diet-induced obesity (DIO) promotes the accumulation of stromal cells with senescent characteristics in the adipose tissue (AT). Selectively clearing these cells—either through chemical senolytics or activation of invariant natural killer T (iNKT) cells—improves glucose homeostasis in obese mice, however the identity of the responsible stromal population remains unknown. Here, we use transcriptional profiling of AT stromal cells coupled with C12FDG-based enrichment of senescent populations in mice with DIO and healthy controls to identify a distinct subset of adipose progenitor cells (APCs) with robust senescence signatures that accumulate in DIO across multiple AT depots. We show that these cells, which we term senescent APCs (sAPCs), are not merely passive markers of metabolic stress but are instead active stromal organizers, accumulating in parallel with the emergence of lipid-associated macrophages (LAMs) and the diminution of multipotent mesenchymal progenitors. sAPCs promoted CCR2-dependent macrophage chemotaxis, directly linking stromal senescence to chemokine-mediated remodeling of the AT immune niche. Comparative transcriptional analysis revealed a remarkable similarity between sAPCs and inflammatory cancer-associated fibroblasts (iCAFs), including the strong induction of periostin (POSTN) and the production of osteoprotegerin (OPG), a decoy receptor for RANKL and TRAIL that enables tumoral immune evasion. Indeed, OPG production by AT stromal cells was induced by DIO across AT depots. Exogenous OPG inhibited the ability of iNKT cells to kill senescent APCs in vitro, whereas antibody-mediated OPG neutralization reciprocally enhanced such cytotoxic killing. In vivo, systemic OPG neutralization both reduced sAPC accumulation in AT and normalized glucose homeostasis in obese mice. Together, these findings identify sAPCs as a pathological stromal population that expands in obesity through elaboration of immunomodulatory factors. In particular, secreted OPG enables sAPCs to evade iNKT-mediated immune surveillance and contributes to metabolic dysfunction, highlighting OPG and sAPCs as promising therapeutic targets for restoring AT immune and metabolic homeostasis.

## INTRODUCTION

Cellular senescence, a state in which cells permanently exit the cell cycle and become resistant to apoptosis, occurs in response to stressors like DNA damage, endoplasmic reticulum stress, and telomere attrition. Senescent cells accumulate in tissues in the setting of advanced age, adopting a senescence-associated secretory phenotype (SASP) that promotes pathological inflammation and age-associated disease manifestations such as musculoskeletal frailty^1,2^.

In adipose tissues (AT), chronic overnutrition and resultant diet-induced obesity (DIO) encourage an enrichment of pathological cell types that resemble senescent cells found in other contexts. We and other groups showed that the build-up of such pathological cells in the AT is a driver of metabolic disruption^3,4^ and cardiovascular disease^5,6^.

Whereas prior studies suggest that senescent cells accumulating in the AT in response to DIO arise from the pool of tissue-resident stromal progenitors^3,7–9^, this work has faced important limitations. Firstly, AT progenitors are highly heterogeneous, with the number of subsets varying based on the depot, age, and state of metabolic health^10^. Examples include early-stage multipotent mesenchymal progenitors with the capacity to differentiate not only into adipocytes, but also fibroblasts, chondrocytes, and osteocytes, as well as later-stage progenitors more specifically committed to the adipocyte lineage, also known as adipose progenitor cells (APCs)^11^. However, we do not know the precise progenitor subset(s) that shift toward senescence with either age or DIO, nor the extent to which such cells in the AT transcriptionally resemble cell types that undergo senescence in other contexts. This is because until now, we have lacked definitive markers to specifically and accurately track and analyze these cells within AT.

Second, new chemical and immunologic strategies are now established to effectively clear senescent cells from tissues including AT, resulting in a restoration of metabolic health in the face of DIO^3,7–9^. For instance, we previously showed that cells exhibiting SASP accumulate in AT due to impaired immune surveillance and clearance by invariant natural killer T cells (iNKT cells), the number and functionality of which are diminished in the context of obesity^12^. By strongly activating iNKT cells in vivo through systemic administration of the CD1d glycolipid antigen alpha-galactosylceramide (αGalCer), we could effectively deplete senescent stromal cells from the AT of mice with DIO and in so doing restore normal glucose homeostasis without inducing weight loss^3^. However, the lack of definitive markers has also precluded us from knowing precisely which AT progenitors, when ablated, are sufficient to improve glucose metabolism in this setting.

Current methods rely on detecting multiple hallmarks, including senescence-associated beta-gal (SA-βgal), and the cell cycle inhibitors Cdkn2a (p16^INK4a^), and p21^WAF1^. However, each of these markers may also be expressed by post mitotic and non-senescent cells^13^. Thus, an unbiased transcriptional examination of the full gamut of AT progenitors is needed to uncover precise markers and/or comprehensive gene sets that accurately define the shifts occurring among progenitor populations, and specifically among APCs, in the setting of DIO.

Here, we leverage recent advances in single-cell RNA sequencing (scRNA-seq) that have been instrumental in elucidating the biological importance of discrete cell populations within complex mammalian tissues^14^ to investigate the cellular composition of the AT stromal compartment in healthy mice and those with chronic DIO, with an eye on identifying unique and mechanistically important senescence markers. Our results reveal the accumulation of a specific senescent, pro-fibrotic, pro-inflammatory population of APCs along with parallel shifts in key aspects of immune cell heterogeneity. These senescent APCs are characterized by increased expression of previously unrecognized genes, including those encoding periostin (POSTN) and the soluble RANK/TRAIL ligand decoy receptor osteoprotegerin (OPG).

We show that the pathological APCs numerically enriched in the AT during DIO secrete high levels of OPG, which allows them to evade clearance by iNKT cells. Indeed, neutralizing extracellular OPG using a specific antibody was sufficient to both limit the build-up of pathological APCs in the AT and restore normal glucose homeostasis during DIO.

## MATERIALS AND METHODS

### Mice

All mouse studies were performed on male C57BL/6J according to the IACUC standards following ethics approval by the animal committee at the University of California San Francisco (UCSF). Dietary studies involved age-matched cohorts of mice that were either fed a high-fat diet (HFD) with 60% kcal from fat (cat. no 380050) or a standard low-fat control chow diet (CD) containing 10% kcal from fat (cat. no 380056) from 6 to 22-24 weeks of age (16-18-week diet duration) prior to being purchased from the Jackson Laboratories (Bar Harbor, ME), or that were first purchased (cat. no 000664) and then similarly fed either the HFD or the CD at UCSF. Mice were housed in a specific pathogen-free facility with a 12-hour light-dark cycle at 23 C.

### Isolation of AT SVF cells

SVF cells were isolated from the epididymal white AT (eWAT) of mice by dissecting epididymal fat pads from euthanized mice, washing them twice in D-PBS, mincing them into a slurry, and then digesting them with 1 U/ml Collagenase-D (Roche) in 1X Hank’s Buffered Saline Solution with Ca++/Mg++ containing 1% BSA (Sigma-Aldrich) for 30min (37° C) with agitation. The eWAT lysate was strained twice through a 100 μm mesh into 30 mL of complete culture medium (DMEM-F12 1:1, 10% FBS, 1X Penicillin-streptomycin), followed by centrifugation at 500 x g for 5 minutes (4° C), brief vortexing to resuspend the SVF pellet, and then spinning again with the pelleted cells decanted & resuspended in ACK lysis buffer to remove contaminating red blood cells. The lysis reaction was neutralized by diluting with D-PBS, centrifuging, decanting, and then resuspending. Live SVF cells were counted using trypan blue and then used for various assays.

### Cells

Mouse iNKT hybridoma DN32.D3 cells were kindly provided by Dr. Mitchell Kronenberg (La Jolla Institute for Allergy and Immunology)^15^ and maintained in RPMI-1640 medium supplemented with 50 μM β-Mercaptoethanol and 10% FBS. SVF cells were isolated as described above and cultured in MEM-α medium supplemented with 10% FBS and 1% Antibiotic-Antimycotic (anti-anti). Bone marrow-derived macrophages (BMDMs) and Bone marrow dendritic cells (BMDCs) were isolated from bone marrow of 6-8 weeks old C57BL/6J mice. BMDMs were cultured in macrophage differentiation medium plus M-CSF/CSF-1 for 6-7 days, and BMDCs were cultured in presence of GM-CSF for 6 days, per published protocols^16^.

### Flow cytometry

eWAT SVF cells were prepared as above, washed in Cell staining Buffer (BioLegend) and stained on ice unless otherwise indicated. Data were acquired on Attune™ NxT Flow Cytometer or Cytek Aurora. Data were analyzed using FlowJo (v10).

#### C12FDG staining

C12FDG staining was performed as previously described^3^. Briefly, eWAT SVF cells were adjusted to 3×10^6^ cells/mL in complete culture medium, incubated with 33 μM C_12_FDG (MarkerGene) for 1 hour (37° C), and then centrifuged at 500 x g for 5 minutes (4° C), washed with Cell Staining buffer (BioLegend), and stained with anti-CD45-APC (clone 30F-11) and anti-CD31-PE-Cy7 (clone 390) for 30 minutes on ice. Cells were washed in Cell Staining buffer and resuspended in Cell Staining buffer containing DAPI or 7-AAD (Cayman Chemical, 400201). C12FDG fluorescence was quantified in live singlet CD45^−^ cells or CD45^−^CD31^−^ cells.

#### iNKT staining

SVF cells were stained with CD1d-PE tetramer (NIH Tetramer Core Facility) loaded with the αGalcer analog PBS57, and with anti-CD3-APC (clone 17A2, Biolegend). Unloaded CD1d-PE tetramer-stained samples were used to determine background staining. Live cells were identified with DAPI, and iNKT cells were further defined as CD3^Low^ and Cd1d Tetramer^+^.

#### Immune cell analysis

For macrophage characterization, SVF cells were stained with anti-CD45-FITC, anit-CD11b-PECy7, anit-CD9-PE, and anti-CD163-APC. CD9^+^ and CD163^+^ populations were gated within CD45^+^CD11b^+^ populations.

### CD45 magnetic separation of SVF cells

Immune cells from eWAT SVF were separated from other stromal cells by anti-CD45 antibody selection using MojoSort magnetic conjugated nanobeads (BioLegend) per manufacturer’s protocol. The eWAT SVF cells were first washed with MojoSort buffer and incubated with 10 mL of mouse anti-CD45-nanobeads for 15 mins on ice. Cells were then washed with MojoSort buffer, centrifuged at 300 x g for 5 minutes (4° C), decanted & resuspended in 3 mL MojoSort buffer. Nanobead-bound cells were placed in a magnetic column for 15 minutes (4° C). The immune cells (CD45-bound, magnetic) and other SVF cells (CD45-unbound, non-magnetic supernatant) were independently collected, centrifuged at 500 x g for 5 minutes (4°C), and then processed for cell culture studies.

### RNA isolation, cDNA synthesis, and real-time qRT-PCR

RNA was extracted with TRIzol reagent using the Zymogen total RNA micro-prep kit (Zymogen) according to manufacturer’s protocol, and total RNA was quantified by Nanodrop. ∼200–300 ng of total RNA was used for cDNA synthesis (qScript cDNA synthesis kit; QuantaBio). cDNA (250 ng) was diluted 1:5 before amplification with Power-UP SYBR reagent using the ABI 384-well QuantStudio 5 Real-time PCR system or the Bio-Rad 96-well quantitative PCR platform. Quantification was performed using the delta-delta C_T_ method, with cyclophilin A (*Ppia*) as a housekeeping gene.

### Chemotaxis Assay

Chemotaxis experiments were conducted using 6-well plates equipped with 5 μm polycarbonate membrane transwell inserts, following established methods^17^. Briefly, eWAT SVF cells (including APC fractions) were isolated from eWAT as described above and depleted of CD45+ immune cell fraction using MojoSort mouse CD45 magnetic nanobeads (Biolegend). Cells were plated (∼300,000 cells/well) and allowed to recover in complete medium for 2 days, and then then treated with 50 mM etoposide for 48 hours to induce senescence. After etoposide washout, cells were incubated in serum-free MEM medium for another 2 days to generate conditioned media. The senescence-conditioned media was collected and placed to the lower chamber of the transwell system.

Separately, primary mouse BMDMs were isolated from the bone marrow of C57BL6/J mice as described^18^ and added to the upper chamber insert. Where indicated, the CCR2 inhibitor RS102895 (10μM) was included during the assay, and recombinant CCL2 (10ng/ml) was used as a positive control, After incubation at 37°C for 6 hours, BMDM that migrated to the lower chamber were counted.

### Measurement of POSTN and OPG by ELISA

The eWAT SVF from CD or HFD-fed mice were isolated as mentioned above and incubated 24-48 hours at 37°C. The standard culture medium was then replaced with a serum-free medium, and the cells were incubated for another 24-48 hours to produce conditioned media, from which POSTN (cat #MOSF20] and OPG (cat #MOP00) levels were measured by ELISA, per manufacturer’s instructions.

### Cytotoxicity assay

Epididymal AT (eWAT) stromal cells were purified by CD45-depletion using MojoSort mouse CD45 nanobeads. Senescence of eWAT stromal cells was induced by treatment with 20 μM etoposide for 24 hrs. BMDC were isolated from male C57BL6/J mice and cultured as described above, after which they were pulsed with either αGalCer (10 or 100 ng/L) or vehicle overnight and then washed with PBS before co-culturing with iNKT hybridoma cells at a 4:1 ratio. To test whether OPG impacts the ability of activated iNKT to kill senescent target cells, etoposide-treated eWAT stromal cells were pretreated with either recombinant OPG or varying concentrations of αOPG, respectively, and then incubated in the presence of α-GalCer-activated iNKT hybridoma cells for 8 hours. The viability of eWAT stromal cells was then measured using the ATPlite Luminescence Assay.

### Glucose tolerance test

Glucose homeostasis was assessed via an intraperitoneal glucose tolerance test (IPGTT) as described previously^3,17^. Briefly, male C57BL6J mice fed the HFD to produce DIO were treated with either 1) 100 mg/kg of the chemical senolytic agent ABT-737 in 30% propylene glycol, 5% Tween-80 and 65% D5W by oral gavage, 2) an established polyclonal anti-αOPG antibody (R&D Systems Catalog #: AF459)^19^, 3) an isotype control antibody (R&D Systems Catalog #: AB-108-C), or appropriate vehicle every week for four weeks. Age- and sex-matched mice fed CD and treated with vehicle served as negative controls. Following treatments, mice were fasted overnight for 14-hours and then injected intraperitoneally with glucose (2 g/kg body weight). Blood glucose was measured by tail vein at 0, 15, 30, 60, and 120 minutes later using a glucometer (Abbott Diabetes Care, Inc., Alameda, CA).

### Single-cell RNA sequencing (scRNA-seq) and data analysis

Single-cell suspensions from eWAT SVF pooled from CD- and HFD-fed mice were stained with C12FDGF as described above and then incubated with Fc block (anti-CD16/32; UCSF Antibody Core) and anti-CD45 (clone 30-F11) for 30 min on ice. Following staining, cells were sorted on a BD FACSAria II to yield immune cells (CD45⁺) and a spectrum of non-immune stromal cells based on C12FDG fluorescence (CD45-C12FDG+). The sorted cells were assessed for viability using a hemacytometer and processed for high-throughput droplet-based scRNA-seq using the 10x Chromium 3’ end platform (10X Genomics). Pseudoalignment with kallisto | bustools was used to generate a gene count matrix^20–22^. Initial preprocessing, including removing empty droplets and quality control using Scanpy (version 1.9.1), was performed^23^.

#### Quality control and normalization

Cells with less than 1000 detected counts and for which the total mitochondrial proportion exceeded 5% were excluded from further analysis. Doublets were then removed using scrublet with default parameters^24^. Genes with a minimum total count of 10 across all cells were also filtered out. Raw counts were normalized to 10,000 counts/cells and were log1p-transformed. For gene expression counts, the top 5000 highly variable genes were selected using the Scanpy function *sc.pp.highly_variable_genes*, employing Seurat flavor with default parameters^25^. Subsequently, an scVI model was trained on the raw counts using all cells^26^. Clustering was performed using *Leiden* algorithm^27^ and visualized using UMAP using default parameters^28,29^.

#### Data analysis on adipose tissue (AT) dataset

Each library (Chow/CD45^−^C_12_FDG^+^, HFD/CD45^−^ C_12_FDG^+^, Chow/CD45^+^; and HFD/CD45^+^) yielded high quality data, resulting in total 36,387 cells. The initial data was split into two datasets based on *Ptprc* expression: The "immune dataset" included *Ptprc* expressing cells (CD45+) and the “stromal dataset” included *Ptprc* non-expressing cells enriched for senescence (CD45-C12FDG+). The raw counts were reprocessed with scVI (version 0.19.0) separately for the stromal and immune datasets using the same workflow as described above and visualized using UMAP with default parameters^26^. Leiden clustering of the immune dataset was performed in Scanpy at a resolution of 0.5, resulting in 15 clusters. Similarly, clustering of the stromal dataset at a resolution of 0.5 initially identified 8 clusters, but some clusters lacked specific markers. Therefore, we reduced the resolution to 0.25, which refined the analysis and resulted in 5 well-defined clusters. One cluster was annotated as endothelial cells based on high *Pecam1* expression and was removed prior to downstream analyses to focus on non-endothelial stromal populations.

#### Data analysis of integrated AT and tumor microenvironment (TME) stromal datasets

Single cell stromal datasets from AT and TME were integrated using scVI^26^. Two AnnData objects containing filtered stromal cells (“AT stromal” and “cancer stromal”) were loaded and concatenated into a single object with unique cell identifiers. The TME stromal dataset comprised GFP^−^ CD45^−^ cells (TME stromal cells depleted of GFP^+^ tumor cells and immune cells). To retain prior annotations, previously processed/annotated versions of each dataset were loaded, and cluster labels were transferred to the concatenated object by matching cell barcodes, including “AT stromal” labels and “cancer stromal” labels.

Preprocessing was performed as described above. Briefly, low-abundance genes were filtered (minimum total counts = 10, minimums cells =10) and raw counts were normalized to 10,000 counts/cells and were log1p-transformed. Highly variable genes were selected (5,000 genes, Seurat v3 method) using sample as the batch key to account for batch structure. A single scVI model was trained on the combined dataset using the raw count layer, with sample specified as the batch variable to correct for technical/sample specific effects. Leiden clustering of the immune dataset was performed in Scanpy resulting in 8 clusters. UMAP was visualized colored by sample/dataset identity and by transferred AT and TME cluster annotations to enable cross-tissue comparison of stromal cell states.

#### Cosine similarity analysis

To quantify how closely AT stromal cells resembled tumor iCAFs, cosine similarity was computed between each AT stromal cell’s scVI-normalized expression profile and an iCAF reference profile. The iCAF reference vector was defined as the mean scVI-normalized expression across cells annotated as cancer cluster 3 (iCAFs) in the cancer stromal dataset. Cosine similarity was then calculated for each AT stromal cell (from the stromal AT dataset) against this iCAF mean vector using *sklearn.metrics.pairwise.cosine_similarity,* using the shared, gene-aligned feature space^30^. The resulting similarity score was added as metadata (“cosine similarity”) to the AT AnnData object by matching cell barcodes and was visualized on UMAP and summarized across stromal clusters using violin plots.

#### Differential gene expression

Differential expression analysis across different clusters or between different treatment groups in the same cluster was performed using Scanpy (*sc.tl.rank_genes_groups*) with the Wilcoxon rank-sum test. Genes with an adjusted p-value < 0.001 and log2 fold change greater or equal to 1 were selected as marker genes.

#### UMAP differential density estimation

A Kernel density estimation approach was used to visualize variations in cell subpopulation density between CD and HFD samples. We separately calculated the densities of each sample using the UMAP coordinates of the respective cells and then extrapolated the densities to the entire two-dimensional UMAP space using the nonparametric kernel_density.KDEMultivariate() function from the statsmodels Python package. At each point in the UMAP space, we calculated and displayed the logarithm of the estimated densities ratio, log (HFD_density/ Chow_density), as a proxy for the likelihood of a UMAP region being associated with an increase (positive log density ratio) or decrease (negative log density ratio) in the proportion of HFD cells^30^.

### Statistics

The number of biological replicates (n) for each study is indicated in the corresponding figure legend. Statistical comparisons were performed between groups of mice, and/or samples (n ≥ 3 biological replicates) using two-tailed t-tests for comparisons between two groups, or either one-way or two-way ANOVA as appropriate for analyses involving three or more groups, with corrections for multiple testing. Significance was defined as p ≤ 0.05. Statistical analyses were performed and associated figures generated using GraphPad Prism 10 software. Biorender software was used to produce illustrations that accompany specific figures.

## RESULTS

### A subset of senescent APCs specifically accumulates in the AT of mice with DIO

Previous studies showed that DIO induces the build-up of APCs in the AT that exhibit features of SASP^3,7–9^. We further showed that stimulating iNKT cells to eliminate such cells from the AT is sufficient to preserve normal glucose homeostasis in mice with DIO^3^. Here, we performed scRNA-seq using the 10X Genomics platform to identify and characterize the specific senescent APC populations that accumulate in AT during DIO.

To do so, we first isolated the SVF from the eWAT of mice fed either a standard low-fat chow diet (CD) or a Western-type high-fat diet (HFD) for 16 weeks (Figure 1A). These SVF cells contained all eWAT cell types other than mature adipocytes. Because beta-galactosidase (β-gal) expression ranges widely across APCs but only correlates with bona fide senescence/SASP markers in the subset of APCs expressing β-gal at the peak of this range^3^, we stained eWAT SVF cells with C_12_FDG (a β-gal reporter) to enrich for such senescent APCs, which we hypothesized might be relatively rare. We next used FACS to separate and isolate both immune cells (CD45^+^) and the non-immune SVF cells that we had enriched for APCs with bona fide senescent features (CD45^−^C12FDG^+^). We processed both fractions for scRNA-seq and performed downstream bioinformatic analysis with scVI^21^.

**Figure 1.**
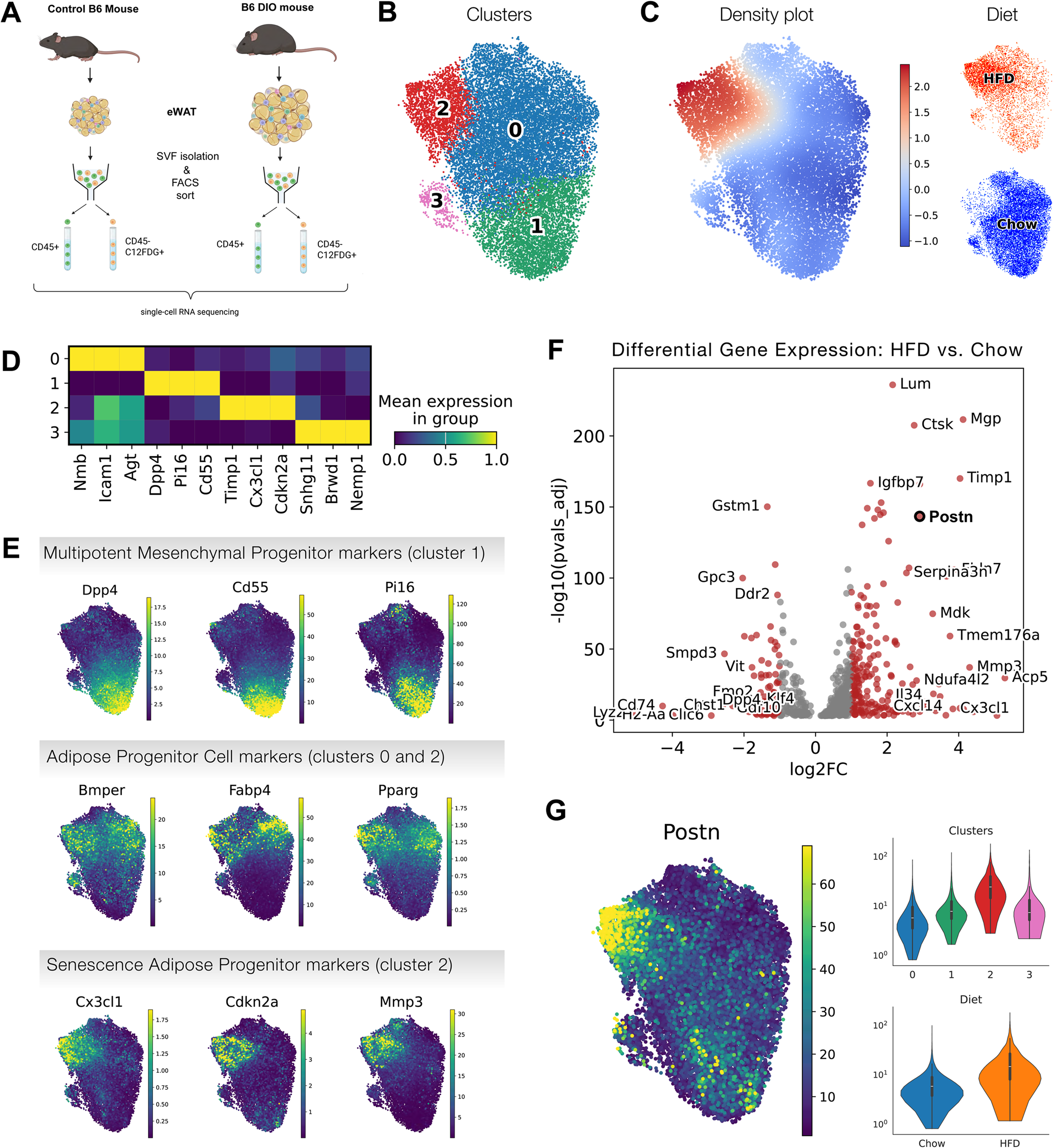
sAPCs accumulate in the eAT of mice with DIO. **(A)** Experimental workflow. **(B)** UMAP feature plot of different clusters in CD45^−^ C12FDG^+^ SVF cells. Distinct clusters are identified using the Leiden algorithm and are represented by different colors. **(C)** UMAP representation of scRNA-Seq analysis of CD45^−^ C12FDG^+^ SVF cells from mice chronically fed either the CD (blue) or HFD (orange), with corresponding density plot showing the difference in density between the different diet groups (positive log density ratio shows HFD-enriched regions in red, and negative log ratio shows CD-enriched regions in blue) **(D)** Matrixplot showing normalized expression of selected genes (cluster characterization) **(E)** UMAP feature plots showing gene expression of selected markers for multipotent mesenchymal progenitor cells, APCs, and sAPCs, respectively **(F)** Differential gene expression analysis of non-immune (CD45^−^ C12FDG^+^) AT stromal cells across both CD and HFD conditions, with volcano plot specifically showing transcriptionally up- and downregulated genes in mice fed the HFD. **(G)** UMAP feature plots and violin plots showing *Postn* expression across AT SVF clusters and between CD- and HFD-fed mice, with values in each plot reflecting scVI normalized counts.

Focusing first on the non-immune (CD45^−^C12FDG^+^) SVF cells, we used unsupervised clustering to identify four transcriptionally distinct clusters after batch effect correction (Figure 1B) that further segregated in transcriptional space based on diet, with CD-derived cells enriched in clusters 0, 1, and 3, and HFD-derived cells predominantly found in cluster 2 (Figure 1C).

Expression of selected marker genes distinguished these clusters (Figure 1D), consistent with previously described AT stromal populations^11^. Cluster 1 was characterized by increased expression of *Dpp4, Pi16*, and *Cd55* (Figures 1D-E), genes known to be expressed by multipotent mesenchymal progenitor cells capable of differentiating not only into adipocytes, but also fibroblasts, chondrocytes, and osteocytes^11^, whereas clusters 0 and 2 were enriched for genes known to be expressed in adipose progenitor cells (APCs) committed to the adipocyte lineage^11^ (Figure 1E).

Although cells in cluster 2 shared several markers with other APC populations, their overall transcriptional landscape was distinct from the other APC subsets, reflecting a state that had not been described in prior studies. In particular and in contrast to cluster 0 cells, which also showed a classical adipogenic lineage-committed APC profile, cells in cluster 2 uniquely expressed atypical cell cycle regulators including cyclin dependent kinase inhibitor 2A (*Cdkn2a*; *p16*), as well as pro-inflammatory mediators such as chemoattractant C-X3-C motif chemokine ligand 1 (*Cx3cl1*) and extracellular matrix remodeling factors such as matrix metallopeptidase 3 (*Mmp3*), which are well known to be pathologically expressed in the context of senescence/SASP (Figure 1E) ^31,32^. Importantly, whereas cluster 0 APCs were derived largely from CD-fed mice, cluster 2 APCs, which we term pathological senescent-APCs (sAPCs), were almost exclusively derived from HFD-fed mice (Figure 1C). This enrichment together with the senescence/SASP gene expression profile that defined cluster 2 (Figure 1E) provides strong evidence for the accumulation of a specific senescent APC population in the AT of mice with diet-induced obesity (DIO).

Using differential gene expression analysis we saw that the transcriptional signature of non-immune AT SVF cells from CD-fed mice aligned with strong representation by cells expressing a multipotent mesenchymal progenitor-like program, marked by genes such as *Dpp4*, *Ddr2*, *Wnt10b* and *Bmp7*^11,33,34^ (Figure S1), whereas such cells were relatively fewer in number in the AT of HFD-fed mice. By contrast, our analysis showed that cells marked by the expression of genes involved in ECM remodeling (*Timp1, Mmp3, Lum*), inflammatory and chemokine signaling (*Ccl8, Cx3cl1, Cxcl9, Il34, Apoe*), and metabolic responses to hypoxic stress (*Ndufa4l2, Cox8b*) were relatively enriched within the non-immune AT stromal compartment of HFD-fed mice when compared with CD-fed controls (Figure 1F)^35–37^.

Notably, the gene encoding periostin (*Postn*), a secreted matricellular protein that is normally expressed during embryonic development but that is lowly expressed or absent in adult postnatal tissues^38,39^ was among the top 10 most upregulated genes in non-immune AT SVF cells from HFD-fed vs. CD-fed mice (Figure 1F). Moreover, *Postn* expression was specifically enriched in sAPCs (cluster 2) when compared against other UMAP clusters. Together, these findings indicate that the sAPCs accumulating in the AT of mice with DIO are marked by upregulation of *Postn*.

### sAPCs in the AT are specifically pro-fibrotic

Senescent cells are implicated in driving fibrosis in multiple tissue and disease contexts^40,41^. Given this and the fact that AT fibrosis is linked to obesity-induced insulin resistance^36^, we examined the fibrotic potential of sAPCs accumulating in the eWAT of HFD fed mice. To do so, we computed a fibrosis-associated gene-set score from a curated panel of fibrosis-related genes and compared this score across the four transcriptionally distinct stromal progenitor clusters and between the two dietary conditions. This analysis revealed that sAPCs (cluster 2 cells) expressed a strong pro-fibrotic gene program when compared to other stromal clusters and that fibrosis scores were overall higher in SVF cells from mice with DIO vs. controls (Figure 2A-C). Consistent with the gene set-score analysis, we found for essentially each gene in the fibrosis-related panel, sAPCs had both a relatively high average expression level as well as an increased fraction of cells expressing pro-fibrotic genes when compared against cells from each of the other clusters (Figure 2C). In particular, most ECM remodeling and fibroblast activation genes were expressed by a relatively small fraction of cells in other clusters but were detected in a comparatively large fraction of sAPCs, indicating a selective expansion of these specific gene programs in this population (Figures 2C-D). This finding was supported by UMAP feature plots and gene-level comparisons, which further confirmed robust upregulation of specific AT fibrosis-associated genes (*Col6a1*, *Col6a3*, *Lum*) in sAPCs (Figure 2D), along with clear evidenced that these same genes were upregulated in AT non-immune SVF cells by HFD feeding^36,42,43^ (Figure 2E). By contrast, genes negatively associated with fibrosis (e.g. *Klf4, Gdf10, Fmo2*) showed a reciprocal pattern, with reduced expression in sAPCs vs cells from the other clusters and downregulation by HFD vs. CD^44–47^ (Figures 2F-G). Together, these findings show that sAPCs fuel a pro-fibrotic transcriptional program within the stromal compartment of AT in the context of DIO, alongside a relative diminution of the anti-fibrotic program seen in the context of a standard CD.

**Figure 2.**
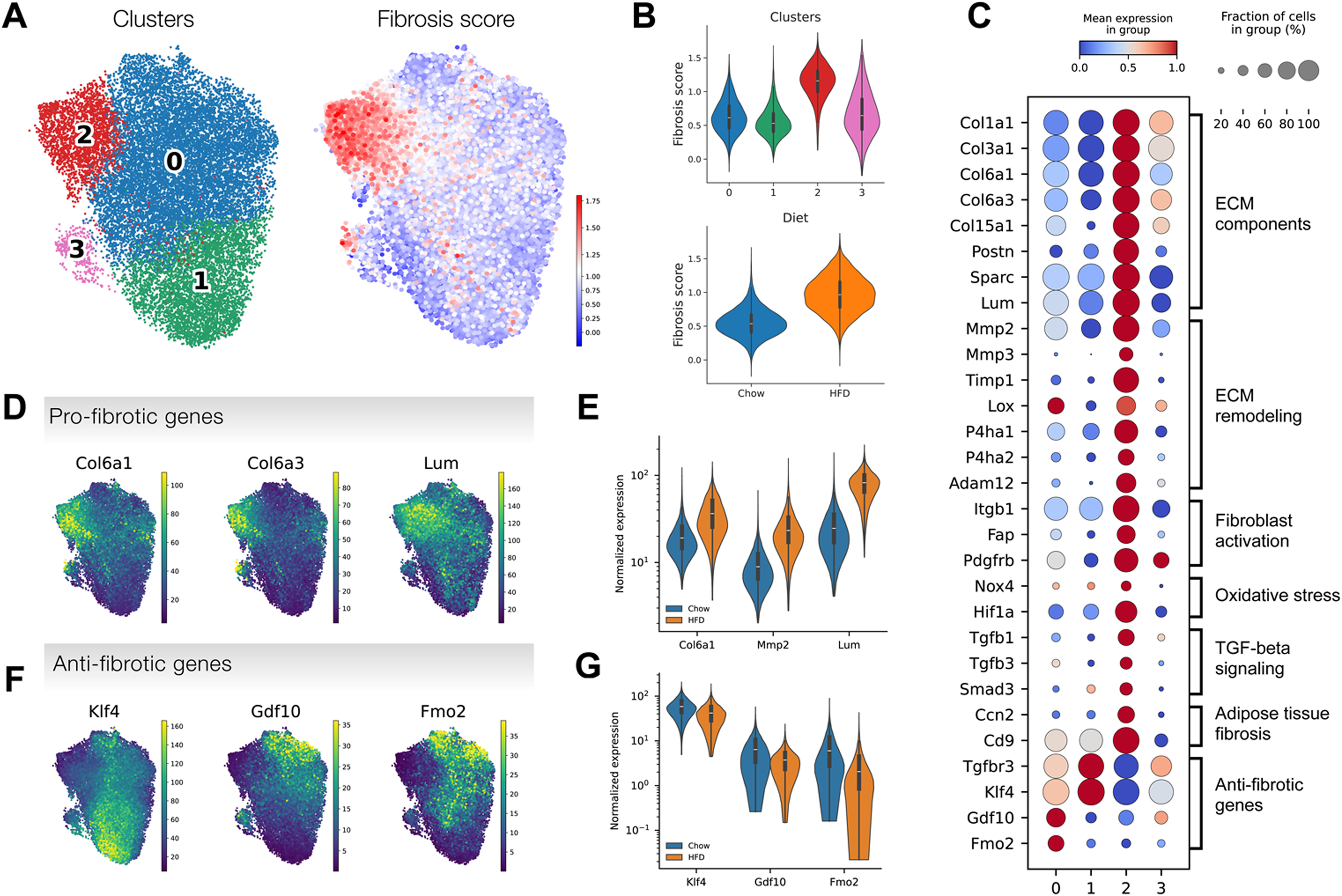
sAPCs express pro-fibrotic genes. **(A)** UMAP feature plots of CD45^−^ C12FDG^+^ SVF cells colored by cluster identity (left) and by fibrosis gene-set score (right). **(B)** Violin plots showing fibrosis score across clusters (top) and between CD- and HFD-fed mice (bottom). **(C)** Dotplot showing the expression of 29 fibrosis-related genes across different clusters. Expression data are scaled between 0 (dark blue) and 1 (dark red). Dot size represents the fraction of cells within each cluster expressing each gene. **(D)** UMAP feature plots showing gene expression of representative pro-fibrotic genes. **(E)** Violin plot comparing expression of representative pro-fibrotic genes between CD- and HFD-derived SVF cells (in scVI normalized counts) **(F)** UMAP feature plots showing expression of representative anti-fibrotic genes **(G)** Violin plot comparing expression of anti-fibrotic genes between CD- and HFD-derived SVF cells. Gene expression values shown in UMAPs and violin plots are shown as scVI normalized counts.

### sAPC buildup in DIO is accompanied by coordinated changes in immune cell composition, suggesting linked stromal–immune remodeling

We next examined changes in the CD45^+^ AT immune cells occurring alongside the sAPC accumulation and the relative loss of multipotent mesenchymal progenitors in the context of chronic DIO. Unsupervised clustering of scRNA-seq data from CD45^+^ eWAT SVF cells identified 16 distinct AT immune populations in the combined pool of CD- and HFD-fed mice (Figures 3A and S2A). Consistent with the literature^48^, the most abundant immune cell populations in the eWAT were macrophages, represented in clusters 0, 1, 3, 7, and 9 (Figures 3A, 3D). Macrophages in clusters 1 and 3 displayed a homeostatic anti-inflammatory program characterized by expression of *Cd163* and *Mrc1* (Figure 3B), whereas those in clusters 0,7 and 9 exhibited a lipid-associated macrophage (LAM) gene signature marked by high expression of lipid receptors and lipid metabolism genes, including *Trem2* and *Fabp4* (Figures 3B and S2B).

**Figure 3.**
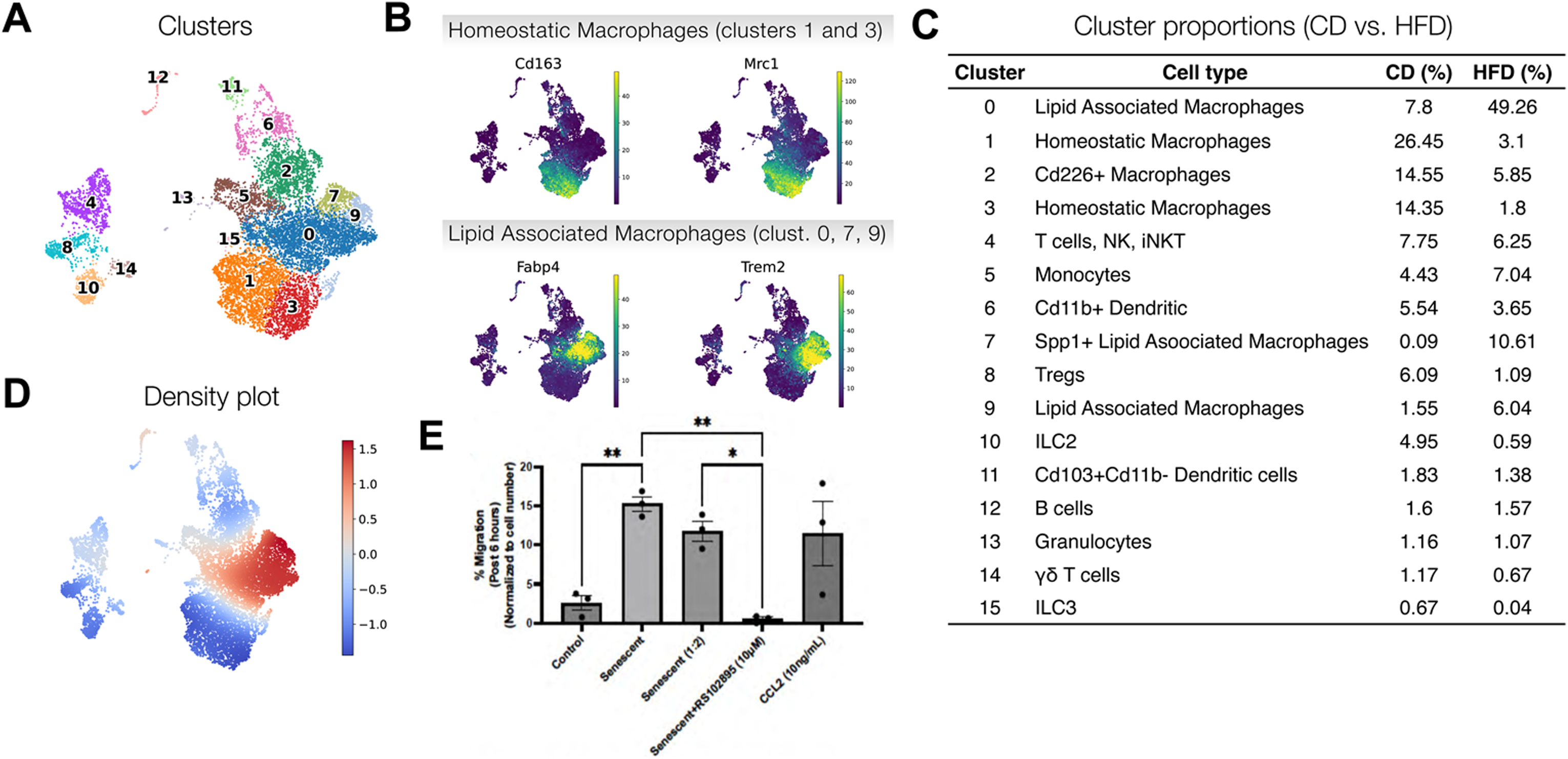
Established shifts in macrophage populations coincide with DIO-associated accumulation of sAPCs in AT. **(A)** UMAP representation of different clusters in AT CD45^+^ cell fractions from mice fed either the CD or HFD. Distinct clusters are identified using the Leiden algorithm and are represented by different colors. **(B)** UMAP feature plot showing gene expression of homeostatic macrophage markers (*Cd163* and *Mrc1) and* Lipid-associated-macrophage (LAM) markers (*Trem2* and *Fabp4*) **(C)** Table showing the percent composition of each cluster in CD vs. HFD. **(D)** Density plot showing the difference in density of AT SVF immune cells derived from HFD- vs. CD-fed mice. Colors show the log density ratio with red showing regions enriched in cells derived from HFD mice and blue showing regions depleted of cells derived from HFD mice. (E) Cell migration assay, showing that medium conditioned by senescent AT progenitors is sufficient to stimulate macrophage recruitment, whereas it is reduced by CCR2 inhibition. Recombinant CCL2 (MCP-1) was a positive control for macrophage chemotaxis. *p<0.05, **p<0.01 by Welch’s t-test.

As expected, comparing macrophage transcriptional subsets by dietary condition showed that chronic HFD consumption induced a pronounced shift away from homeostatic states and towards a LAM signature (Figure 3D) ^49^. Indeed, whereas macrophages in clusters 1 and 3 comprised 26% of eWAT immune cells in lean control mice, they only accounted for 6% in the setting of DIO (Figure 3C). By contrast, whereas LAMs accounted for 8% of eWAT immune cells in CD-fed control mice, their frequency increased to 49% in the eWAT of mice with DIO (Figure 3C). Flow cytometric assessment of eWAT CD11b^+^ myeloid cells in mice over increasing durations of HFD feeding showed a progressive expansion of CD9^+^ LAMs accompanied by a parallel reduction in the frequency of CD163^+^ myeloid cells. Indeed, within 4 weeks of HFD consumption, the frequency of LAMs among CD11b^+^ myeloid cells in the eWAT rose to levels comparable to those of CD163^+^ myeloid cells that predominated in the eWAT of CD-fed control mice, and longer HFD durations caused these LAM numbers to rise even further, greatly overtaking the number of CD163^+^ cells in the process (Figure S2C).

Notably, regulatory T cells (Tregs, cluster 8) and homeostatic innate lymphoid type 2 cells (ILC2s, cluster 10) were also reduced in the eWAT of mice with DIO, from 6% to 1% and 5% to 0.5%, respectively, consistent with changes expected in obesity^50,51^ (Figure 3C). Dendritic cells (clusters 6 & 11), monocytes (cluster 5), T cells + NK cells + iNKT cells (cluster 4), γδ T cells (cluster 14), B cells (cluster 12), granulocytes (cluster 13), and ILC3s (cluster 15) were also identified but with less notable differences between DIO and control (Figure 3C). Too few iNKT cells were sequenced to confirm our previous observations by flow cytometry that iNKT cells are reduced in the AT of mice with DIO^3^. Overall, our scRNA-seq analysis of the immune compartment of eWAT aligns well with the established literature, both validating the quality of our dataset and placing the accumulation of sAPCs and the parallel loss of multipotent mesenchymal progenitors into context with overall shifts in the cellular makeup of AT that occur during DIO.

### sAPCs induce macrophage chemotaxis in a CCR2-dependent manner

We further explored whether sAPCs play a role in driving the build-up of LAMs in the AT of mice with DIO. Using a transwell migration assay, we compared primary bone marrow-derived macrophage (BMDM) chemotaxis towards media conditioned either by eWAT-derived CD45^−^ SVF cells treated with vehicle (control) or those treated with etoposide to induce senescence. Only ∼3% of the BMDMs in this assay migrated toward the medium conditioned by the vehicle-treated eWAT SVF cells, whereas approximately 5-fold more (∼15%) BMDMs migrated towards the medium that had been conditioned by senescent eWAT SVF cells (Figure 3E), indicating that replicative senescence induces non-immune eWAT stromal cells to produce macrophage chemotactic factor(s).

To determine whether this chemotactic response involved CCR2, we added the CCR2 inhibitor RS102895 (10 μM) to the transwell assays above, and found that doing so completely abrogated BMDM chemotaxis toward the medium conditioned by etoposide-treated eWAT SVF cells (Figure 3F). Conversely, BMDMs migrated towards recombinant CCL2 (10 ng/mL) that was added to the lower chamber of the transwell system even in the absence of eWAT SVF conditioned medium, and did so at a rate mirroring what was seen in response to senescence- conditioned medium (Figure 3E). Together, these data indicate that senescent non-immune AT stromal cells signal the chemotactic recruitment of macrophages in a CCL2/CCR2-dependent manner.

### sAPCs transcriptionally resemble inflammatory Cancer-Associated Fibroblasts (iCAFs)

Inflammatory cancer-associated fibroblasts (iCAFs) are a subgroup of CAFs found in solid tumors and are characterized by secretion of inflammatory cytokines and growth factors driven primarily by NF-κB signaling; they are thought to promote tumoral immunosuppression by recruiting myeloid-derived suppressor cells into the tumor microenvironment (TME)^52,53^. Importantly, we recently showed that iCAFs themselves evade immune detection in the TME^38^, a capacity that has also been posited for senescent cells within the AT in obesity^2,3^. We thus asked whether sAPCs emerging in the eWAT of obese mice share transcriptional features with iCAFs.

To enable a direct and unbiased cross-tissue comparison, we examined the combined transcriptional datasets from i) CD45^−^C12FDG^+^ AT stromal cells taken from both CD- and HFD-fed mice and ii) TME of EO771 heterotopic breast cancer stromal cells depleted of both immune cells (CD45^+^) and tumor cells (GFP). We then integrated these two datasets to embed AT and TME stromal cells in a shared space while minimizing sample-associated technical effects. We brought the original AT and CAF cluster annotations into our integrated dataset to track cell-of-origin and compare cluster identities after integration. Joint visualization in a shared UMAP space followed by unsupervised clustering revealed transcriptional convergence between AT sAPCs and iCAFs despite the two cell types arising from distinct pathological and tissue contexts (Figures 4A and S3). Quantification of cosine similarity showed that among AT stromal clusters, sAPCs (AT dataset; cluster 2) had the highest transcriptional similarity to iCAFs, underscoring the specific convergence between obesity-associated sAPCs and iCAFs (Figure 4B).

**Figure 4.**
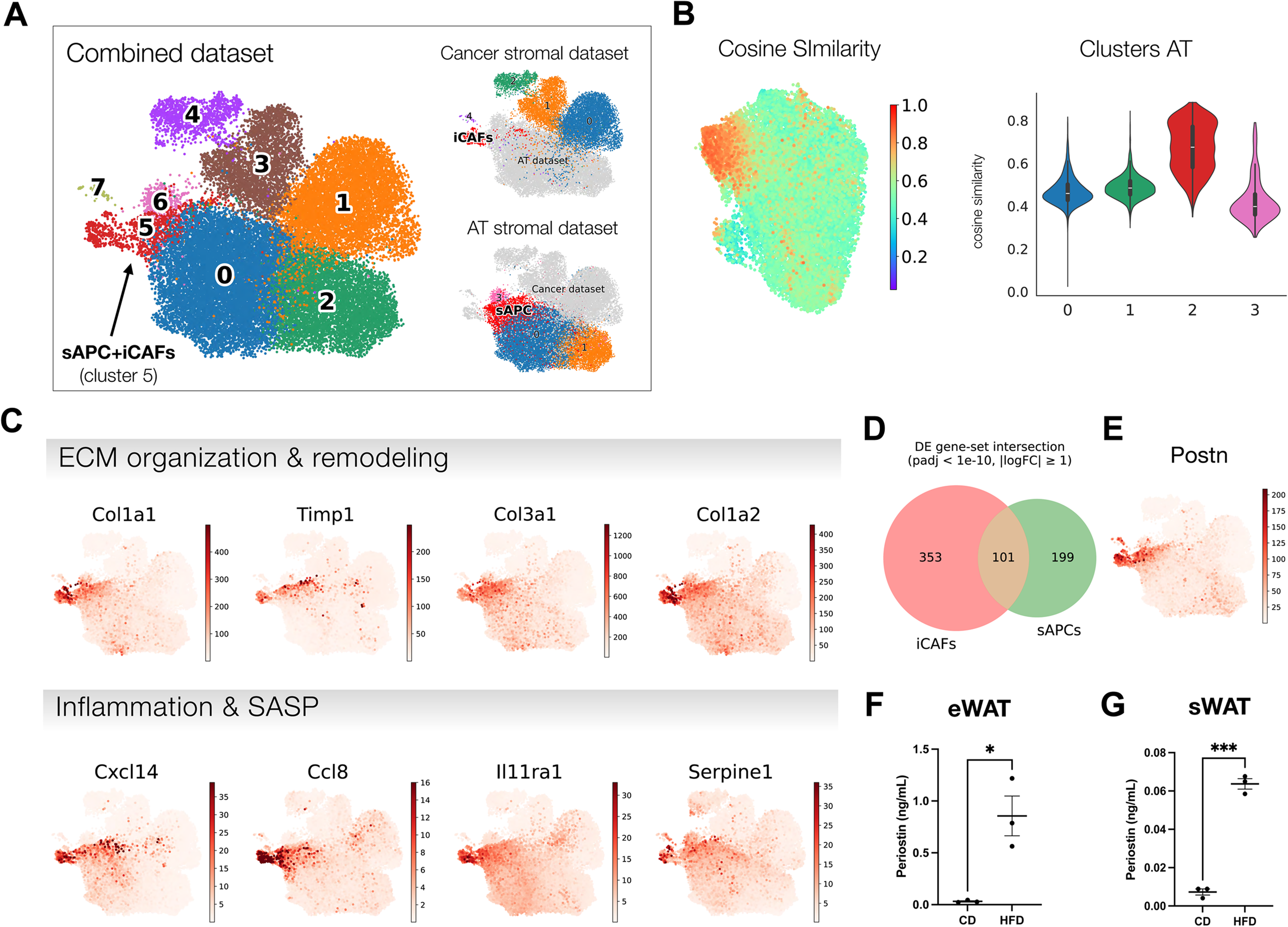
DIO-associated sAPCs transcriptionally resemble tumor-associated iCAFs. **(A)** UMAP visualization of an integrated scRNA-seq atlas combining eWAT stromal cells from CD- and HFD-fed mice with tumor-associated stromal cells^38^. **(B)** Cosine similarity analysis comparing iCAF transcriptional profiles to those eWAT non-immune SVF clusters. **(C**) UMAP feature plots of representative genes commonly expressed by both sAPCs and iCAFs. **(D)** Venn diagram of differentially expressed genes, showing the overlap between iCAFs and sAPCs. **(E)** UMAP feature plot of *Postn* expression in the integrated dataset. **(F-G)** POSTN secretion, as measured by ELISA, from AT non-immune SVF cells, showing that SVF cells from the eWAT (F) and sWAT (G) of DIO mice secrete substantially more POSTN into the medium than do comparable cells from CD-fed control mice. *p<0.05, ***p<0.001 unpaired two-tailed Welch’s t-test.

To identify genes contributing to this transcriptomic similarity, we analyzed the 353 genes that were differentially expressed in iCAFs vs. other clusters in the cancer stromal dataset, and the 199 genes that were differentially expressed in sAPCs vs. other clusters in the AT stromal dataset (Figure 4D). We then assessed the intersection of these two gene sets and identified 101 cluster-defining genes the expression of which was shared by both iCAFs and sAPCs (Figure 4D). These shared genes were enriched for pathways related to ECM organization (*Col3a1, Col1a1, Col1a2*) and remodeling (*Timp1, Mmp3, Mmp14, Adam12*), as well as pro-inflammatory cytokines and chemokines (*Il11ra, Cxcl14, Ccl8, Serpine1*) consistent with an inflammatory, SASP-like transcriptional program (Figures 4C and S3C)^54,55^.

Critically, our integrated analysis also showed that both obesity-associated sAPCs from eWAT and tumor-associated iCAFs exhibit high *Postn* expression exclusively, suggesting that POSTN could serve as an effective handle for fate mapping in both cell types (Figure 4E). Indeed, APC-containing SVF cells from both the eWAT and subcutaneous WAT (sWAT) of mice with DIO secreted nearly ten times more POSTN than did SVF cells from the same AT depots of CD-fed counterparts (Figures 4F-G).

### OPG enables senescent APCs to evade cytotoxic clearance by iNKT cells

Based on the transcriptional similarity between iCAFs and sAPCs, we next sought to determine the extent to which molecular pathways with established functional significance in iCAFs are also engaged in sAPCs. Specifically, our prior work showed that iCAFs secrete osteoprotegerin (OPG), a soluble decoy receptor that binds both RANKL (receptor activator of nuclear factor kappa-Β ligand) and TRAIL (TNF-related apoptosis-inducing ligand), two ligands implicated in T cell and iNKT cell cytotoxic function and thus inhibits their signaling activity^56^. We therefore measured OPG secretion by CD45^−^ AT SVF cells and found that DIO markedly increased OPG secretion by stromal cells isolated from either eWAT or sWAT, when compared with analogous cells from matched AT depots of healthy CD-fed control mice (Figures S4A-B).

We next wondered whether pathological OPG production by sAPCs enables them to evade cytotoxic killing by iNKT cells, effector cells that we previously showed are exhausted in DIO and are no longer able to efficiently clear senescent cells within AT^3^. To probe this question, we established an in vitro system to quantify iNKT-cell activation and cytotoxicity toward an sAPC-like cellular target. First, we validated that αGalCer, a glycolipid antigen that activates iNKT cells through presentation by CD1d^1^, induced mouse splenocyte-derived iNKT cells to express TRAIL, secrete IFNγ, and proliferate (Figure 5A).

**Figure 5.**
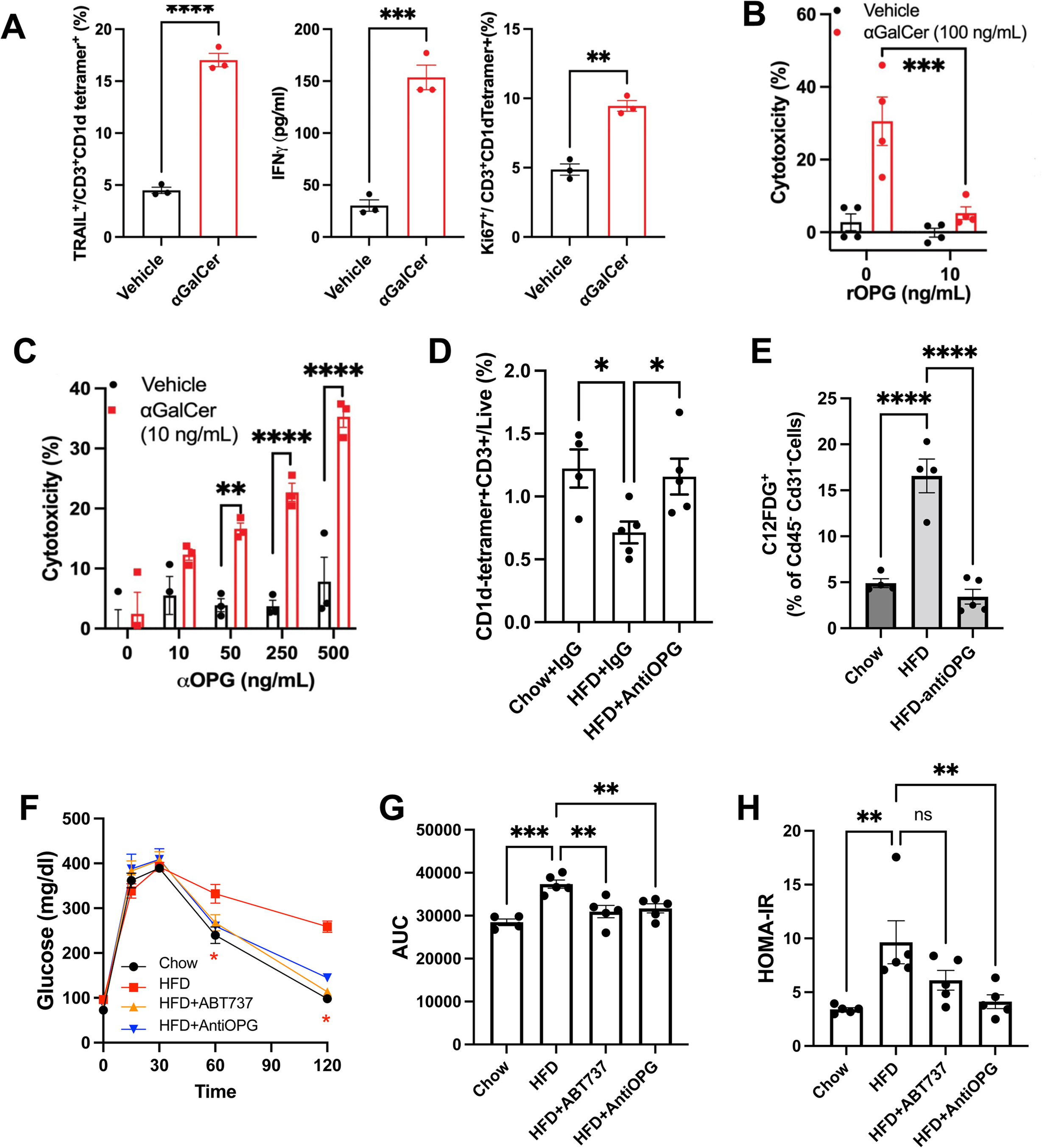
Using an antibody to neutralize OPG reverses glucose intolerance and insulin resistance in mice with DIO. **(A)** Flow cytometry and ELISA data indicating that CD3^+^CD1d tetramer^+^ iNKT cells upregulate TRAIL and IFNγ as they proliferate (Ki67^+^%) in response to αGalCer-dependent activation. **(B)** Cytotoxicity assay, showing that αGalCer-pulsed BMDCs enable iNKT hybridoma cells to kill etoposide-treated adipose progenitor cells (APCs); this capacity is abolished in the presence of recombinant OPG (rOPG, 10ng/mL). **(C)** Anti-OPG antibody (αOPG) dose-dependently rescues the capacity of iNKT hybridoma cells to kill etoposide-treated adipose stromal cells despite being activated by BMDCs pulsed with a low dose of αGalCer (10 ng/ml), which is otherwise insufficient to elicit robust iNKT effector function. **(D)** Systemic αOPG of mice with DIO restores iNKT cell activation in eWAT **(E)** Systemic αOPG treatment of mice with DIO normalizes levels of C12FDG^+^ CD45^−^ CD31^−^ stromal cells to those akin to CD-fed control mice. **(F-H)** Systemic αOPG treatment of mice with DIO normalizes both glucose tolerance (F-G) and the HOMA-IR index of insulin resistance (H). *p<0.05, **p<0.01, ***p<0.001, ****p<0.0001 by ANOVA.

Next, we examined the ability of DN32.D3 mouse iNKT hybridoma cells, cytotoxic effectors known to target senescent cell populations upon αGalCer-dependent stimulation, to kill sAPC-like cells in vitro. We modeled sAPCs by treating mouse AT mesenchymal progenitor cells with etoposide, which we confirmed to responsively secrete OPG into the medium. This enabled us to test the extent to which OPG produced by these sAPC-like cells could modulate the ability of CD1d-activated DN32.D3 iNKT cells to effectively target them for cytotoxic killing. Interestingly, whereas DN32.D3 iNKT cells co-cultured with BMDCs pulsed with vehicle were unable to engage in cytotoxic killing when subsequently exposed to etoposide-treated AT progenitor cells, did not enable them to kill etoposide-treated AT progenitors, DN32.D3 iNKT cells activated by co-culture with BMDCs that had been pulsed with 100 ng/mL α-GalCer became potent cytotoxic effectors that engaged significant killing of sAPC-like senescent progenitors (Figure 5B).

Importantly, the cytotoxic effect of α-GalCer-CD1d-activated DN32.D3 iNKT cells toward their senescent AT progenitor targets was sharply mitigated by adding exogenous recombinant OPG protein to the culture medium in a dose-dependent manner, suggesting that OPG dampens iNKT cell effector-target interaction (Figure 5B). To test whether endogenous OPG secreted by target cells also inhibits the cytotoxic capacity of iNKT cells, we co-cultured DN32.D3 iNKT cells with BMDCs that had been pulsed with only 10 ng/mL of α-GalCer and confirmed that this at this level of stimulation, endogenously secreted OPG could appreciably prevent iNKT cells cytotoxicity of sAPC-like senescent progenitors (Figure 5C). Remarkably, however, adding an OPG neutralizing antibody (αOPG) into the medium in which the DN32.D3 iNKT cells and senescent progenitors were co-cultured was able to restore the cytotoxic capacity of the iNKT cells in a dose-dependent manner (Figure 5C). Together, these data indicate that OPG secreted by senescent AT stromal cells mirroring sAPCs directly inhibits the cytotoxic effector function of CD1d-activated iNKT cells and suggests that the documented inability of activated iNKT cells to efficiently eliminate sAPCs from AT during DIO may be the result of local OPG-mediated suppression of their effector function.

To test this concept in vivo, we systemically administered either αOPG or an IgG control antibody to mice with DIO and examined the impact of each treatment on sAPC content in the eWAT. As compared to control, αOPG treatment restored iNKT cell levels in eWAT, which was suppressed by chronic HFD feeding, to levels similar to those seen in healthy CD-fed mice (Figure 5D). In parallel, αOPG administration completely reversed the HFD-induced build-up of sAPCs in the eWAT (Figure 5E). Together, these findings indicate that systemic neutralization of secreted OPG potentiates iNKT cell activation and alleviates sAPC build-up in the AT during DIO.

### Systemic neutralization of secreted OPG restores glucose homeostasis in mice with DIO

Since systemic αGalCer treatment was shown to restore normal glucose homeostasis in mice with chronic DIO^3^, we therefore next examined whether using systemic αOPG treatment to deplete DIO-associated sAPCs from AT could reproduce this benefit. Remarkably, treating mice that had been chronically fed the HFD with αOPG was sufficient to restore both glucose tolerance and insulin sensitivity, as assessed by the HOMA-IR index, to levels resembling those seen in healthy CD-fed mice (Figures 5F-H). Indeed, these improvements in glucose homeostasis were on par with those achieved by treating mice with the established chemical senolytic drug ABT737, which we used as a positive control (Figures 5F-H). These results indicate that neutralizing secreted OPG to promote the immunologic clearance of sAPCs, a function otherwise impaired in the setting of DIO, is sufficient to normalize glucose homeostasis.

## Discussion

Adipocytes constitute the parenchyma of ATs and are embedded within a connective lattice formed by a highly heterogeneous array of SVF cells. These SVF cells include lineage-committed progenitors that generate adipocytes, fibroblast-like cells that provide structural support, early-stage multipotent mesenchymal progenitors, and a variety of tissue-resident cells that provide immunological support. Together, these cells enable ATs to adapt to a wide range of metabolic stresses. This adaptive capacity is lost during chronic DIO in conjunction with key alterations in the cellular makeup of ATs. Most notably, the immune compartment is known to exhibit a dropout of specific homeostatic macrophage populations, Tregs, and ILC2s, while allowing LAMs to accumulate instead. Our analysis here reproduced these shifts among immune cell populations, a reassuring sign that underscored the high quality of our workflow. Moreover, we show that senescent APCs are capable of chemoattracting macrophages in a CCL2/CCR2-dependent manner.

Here, we show that sAPCs, a unique population of APCs that display a pro-fibrotic transcriptional program and key features of senescence/SASP, accumulate in the ATs of mice specifically in response to chronic DIO. Interestingly, these sAPCs, which emerge in conjunction with a series of DIO-associated shifts in both progenitor and immunologic cell populations, bear a striking transcriptional resemblance to iCAFs observed in TME, including the shared expression of POSTN. iCAFs within TMEs evade detection and clearance by CD8 T cells. Similarly, we find that senescent stromal cells that are normally recognized and eliminated by iNKT cells in adipose tissue escape immune clearance in the setting of diet-induced obesity. Prior studies have identified secreted OPG as a key mechanism by which iCAFs evade immune surveillance. Consistent with this paradigm, we show that OPG plays a comparable role in limiting the cytotoxic activity of activated iNKT cells against sAPCs. These findings indicate that sAPC accumulation in obesity is not a passive consequence of immune dysfunction but instead results from active, OPG-mediated suppression of iNKT cell–dependent immune clearance by sAPCs themselves.

Remarkably, neutralizing secreted OPG using a specific antibody was sufficient to enable activated iNKT cells to effectively kill senescent APCs in vitro, as well as the sAPCs that build up in the ATs of mice with DIO when the antibody was administered in vivo. This systemic neutralization of OPG was importantly also sufficient to normalize glucose homeostasis in the context of DIO, akin to what we previously accomplished by strongly activating iNKT cells using αGalCer^3^. Thus, the insulin resistance and associated glucose intolerance that occurs in mice chronically fed a HFD at least partially reflects the pathological effects of sAPCs that accumulate in ATs through OPG-dependent evasion of immunological surveillance.

We also assessed stromal–immune remodeling in the setting of DIO from the standpoint of the non-immune SVF cells. Importantly, we included staining with C12FDG, a broad indicator of β-gal expression, as part of our workflow in order to allow us to not only detect abundant stromal populations that exhibit low-grade β-gal expression, but also bona fide senescent cells pathologically marked by comparatively high levels of C12FDG staining. Prior scRNA-seq analysis of mouse ATs, including comparisons between relatively lean and obese mice, have not included such C12FDG-based enrichment to search for potentially rare senescent SVF cell populations. As such, these studies may have missed specifically identifying sAPC accumulation in obese ATs, as these relatively rare cells were likely to be lumped together with other APC subsets without the enrichment strategy we employed. Our two-step process prior to hierarchical clustering, by contrast, allowed for separation of the sAPC cluster (cluster 2, only found in the eWAT of mice with DIO) from other APC and progenitor clusters without loss of other non-senescent core stromal populations.

Of note, a recent study used genetic depletion strategies and cellular implantation approaches to implicate senescent endothelial cells in adipose tissue inflammation, metabolic dysfunction, and the loss of glucose tolerance seen in obese mice^57^. These findings are interesting, although it remains unclear precisely how senescence among Tie-2-expressing vascular endothelial cells is linked to metabolic compromise; multiple possible mechanisms, both within tissues and involving inter-organ crosstalk as well as indirect effects stemming from vascular insufficiency could be invoked. It will be interesting to see how future studies delve more deeply into the specific molecular processes underlying this study. Our scRNA-seq analysis included a cluster that was annotated as being endothelial cells based on high *Pecam1* expression. We found that including or excluding this cluster did not impact the transcriptomic data we obtained from any of the other SVF clusters, either immune or non-immune. As such, we removed this cluster prior to downstream analyses to focus on non-endothelial stromal populations. This allowed us not only to more easily identify the sAPCs at the core of this study, but also to assign the molecular mechanism by which sAPCs accumulate to OPG-mediated immune escape, with direct implications on glucose homeostasis in DIO.

The sAPCs we detected exclusively in ATs from mice with DIO shared the expression of certain markers (e.g., *Pparg*, *Fabp4*) with other populations of APCs that were abundant in mice fed the CD and though present, quite numerically diminished in mice with DIO. Expression of these shared markers suggests that the SVF cells we found undergoing senescence were indeed APCs. However, sAPCs emerging in DIO also expressed multiple sets of pro-fibrotic genes as well as other canonical senescence/SASP markers, distinguishing them from the other APC populations. Prior efforts to readily identify senescent cell types within complex tissues have been hampered by the fact that true senescence markers remain poorly defined, forcing investigators to rely on cumbersome composite signatures. By sequencing a substantial number of sAPCs in our analysis, however, we were able to identify POSTN as a robust marker that is highly enriched within the AT specifically in sAPCs in the context of DIO. POSTN is an extracellular matrix protein implicated in tissue remodeling and wound repair. In human WI-38 and IMR-90 fibroblasts, etoposide treatment induced strong POSTN expression alongside canonical senescence markers (*Cdkn2a/p16*, *Mmp2*, *Il8*), and POSTN was secreted into the extracellular milieu, consistent with its inclusion as a factor secreted as part of the SASP (unpublished observations).

Notably, POSTN also marks iCAFs in the tumor microenvironment, a notable fact given that meta-analyzing our scRNA-Seq dataset against that of stromal cells from heterotopic breast tumors^56^ revealed that sAPCs and iCAFs are bear a striking transcriptional similarity. Specifically, both sAPCs and iCAFs express the immunomodulator OPG, which suppresses CD4⁺ and CD8⁺ T-cell activity in tumors and inhibits iNKT cell function within ATs in the setting of obesity. Indeed, both iCAFs and as we show here, sAPCs, secrete OPG.

OPG (TNFRSF11B) is a multifunctional decoy receptor that inhibits two distinct signaling pathways, namely TRAIL-mediated apoptosis, by binding TRAIL and preventing engagement of death receptors, and RANK/RANKL signaling by sequestering RANKL and blocking its interaction with RANK^58^. Both TRAIL and RANKL are induced on activated CD8⁺ T cells, positioning OPG as a potent regulator of T-cell function. By secreting OPG, stromal cells impose a dual brake on T-cell activity that suppresses TRAIL-driven cytotoxicity and neutralizes RANKL-dependent co-stimulatory signals. This establishes an immunosuppressive niche analogous to what is achieved by checkpoint pathways. Indeed, blockade of OPG with a neutralizing antibody restored T-cell effector function, functionally resembling PD-1 or CTLA-4 inhibition. These findings highlight OPG as a stromal-derived immune checkpoint that converges on two fundamental axes of T-cell regulation.

The current study importantly extends this understanding to metabolic tissues, notably ATs. Indeed, OPG emerged as a key secretory factor that mechanistically links the immunosuppressive function of sAPCs in avoiding iNKT cell-mediating cytotoxic clearance with that of iCAFs evading T cell-mediated detection and clearance. This information indicates that metabolic derangements occurring in tissues in the context of chronic DIO may result from the accumulation of pathological, pro-inflammatory and pro-fibrotic cell types that build-up in tissues by altering the balance of stromal-immune interactions in their favor, thus creating a durable niche. In this way, figuring out how to either reengage immunological surveillance mechanisms or “uncloak” potentially pathological stromal cell types to make them easier to clear, could represent emerging strategies to restore healthy metabolic function to complex tissues as well as systemically. As such future studies should probe modulators of such stromal-immune interactions to find new targets. Indeed, we previously highlighted αGalCer-mediated CD1d-dependent stimulation as a means to rev up iNKT cell-mediated senescent cell clearance. Here, we show that neutralizing OPG can “uncloak’ sAPCs, making them easier for iNKT cells to clear in the context of DIO.

The last two decades of research in AT biology has been siloed, focusing either on adipogenic cell types and their lineage commitment and physiological functions, or on resident and infiltrating immune cell types and their role in fomenting “metabolic inflammation” in obesity. Here we show that these two aspects of AT cellular heterogeneity are inextricably linked, and that a specific subset of APCs with senescent features accumulates specifically in the context of DIO by breaking free of normal immunological containment. By doing so, these sAPCs disrupt normal immune-stromal crosstalk to produce inflammatory effects, pro-fibrotic effects, and important impairments in glucose homeostasis.

## Abbreviation List

DIO: diet-induced obesity
SASP: senescence-associated secretory phenotype
iNKT cells: invariant natural killer T cells
AT: adipose tissue
eWAT: epididymal adipose tissue
sWAT: subcutaneous adipose tissue
APCs: adipose progenitor cells
sAPCs: senescent adipose progenitor cells
POSTN: periostin
OPG: osteoprotegerin
RANKL: receptor activator of nuclear factor kappa-Β ligand
TRAIL: TNF-related apoptosis-inducing ligand
αGalCer: alpha-galactosylceramide
scRNA-seq: single-cell RNA sequencing
UCSF: University of California San Francisco
HFD: high-fat diet
CD: chow diet
*Ppia*: cyclophilin A
BMDC: bone marrow-derived dendritic cells
Cdnk2a: cyclin dependent kinase inhibitor 2A
Cx3cl1: C-X3-C motif chemokine ligand 1
Mmp2: matrix metallopeptidase 2
iCAF: inflammatory cancer-associated fibroblast
rOPG: recombinant OPG
αOPG: anti-OPG neutralizing antibody

## REFERENCES

1. Coppé JP, Desprez PY, Krtolica A, Campisi J. The Senescence-Associated Secretory Phenotype: The Dark Side of Tumor Suppression. Annual Review of Pathology: Mechanisms of Disease. 2010;5(1):99–118. doi:10.1146/annurev-pathol-121808-102144

2. Kirkland JL, Tchkonia T. Cellular Senescence: A Translational Perspective. EBioMedicine. 2017;21:21–28. doi:10.1016/j.ebiom.2017.04.013

3. Arora S, Thompson PJ, Wang Y, et al. Invariant natural killer T cells coordinate removal of senescent cells. Med. 2021;2(8):938–950.e8. doi:10.1016/j.medj.2021.04.014

4. Alessio N, Acar MB, Demirsoy IH, et al. Obesity is associated with senescence of mesenchymal stromal cells derived from bone marrow, subcutaneous and visceral fat of young mice. Aging. 2020;12(13):12609–12621. doi:10.18632/aging.103606

5. López-Otín C, Blasco MA, Partridge L, Serrano M, Kroemer G. The Hallmarks of Aging. Cell. 2013;153(6):1194–1217. doi:10.1016/j.cell.2013.05.039

6. Childs BG, Durik M, Baker DJ, van Deursen JM. Cellular senescence in aging and age-related disease: from mechanisms to therapy. Nat Med. 2015;21(12):1424–1435. doi:10.1038/nm.4000

7. Wang L, Wang B, Gasek NS, et al. Targeting p21Cip1 highly expressing cells in adipose tissue alleviates insulin resistance in obesity. Cell Metab. 2022;34(1):75–89.e8. doi:10.1016/j.cmet.2021.11.002

8. Palmer AK, Xu M, Zhu Y, et al. Targeting senescent cells alleviates obesity-induced metabolic dysfunction. Aging Cell. 2019;18(3). doi:10.1111/acel.12950

9. Hickson LJ, Langhi Prata LGP, Bobart SA, et al. Senolytics decrease senescent cells in humans: Preliminary report from a clinical trial of Dasatinib plus Quercetin in individuals with diabetic kidney disease. EBioMedicine. 2019;47:446–456. doi:10.1016/j.ebiom.2019.08.069

10. Emont MP, Jacobs C, Essene AL, et al. A single-cell atlas of human and mouse white adipose tissue. Nature. 2022;603(7903):926–933. doi:10.1038/s41586-022-04518-2

11. Merrick D, Sakers A, Irgebay Z, et al. Identification of a mesenchymal progenitor cell hierarchy in adipose tissue. Science (1979). 2019;364(6438). doi:10.1126/science.aav2501

12. Lee BC, Kim MS, Pae M, et al. Adipose Natural Killer Cells Regulate Adipose Tissue Macrophages to Promote Insulin Resistance in Obesity. Cell Metab. 2016;23(4):685–698. doi:10.1016/j.cmet.2016.03.002

13. Sharpless NE, Sherr CJ. Forging a signature of in vivo senescence. Nat Rev Cancer. 2015;15(7):397–408. doi:10.1038/nrc3960

14. Zheng GXY, Terry JM, Belgrader P, et al. Massively parallel digital transcriptional profiling of single cells. Nat Commun. 2017;8(1):14049. doi:10.1038/ncomms14049

15. Sidobre S, Hammond KJL, Bénazet-Sidobre L, et al. The T cell antigen receptor expressed by Vα14 *i* NKT cells has a unique mode of glycosphingolipid antigen recognition. Proceedings of the National Academy of Sciences. 2004;101(33):12254–12259. doi:10.1073/pnas.0404632101

16. Toda G, Yamauchi T, Kadowaki T, Ueki K. Preparation and culture of bone marrow-derived macrophages from mice for functional analysis. STAR Protoc. 2021;2(1):100246. doi:10.1016/j.xpro.2020.100246

17. Thompson PJ, Shah A, Ntranos V, Van Gool F, Atkinson M, Bhushan A. Targeted Elimination of Senescent Beta Cells Prevents Type 1 Diabetes. Cell Metab. 2019;29(5):1045–1060.e10. doi:10.1016/j.cmet.2019.01.021

18. Gonçalves R, Kaliff Teófilo Murta G, Aparecida de Souza I, Mosser DM. Isolation and Culture of Bone Marrow-Derived Macrophages from Mice. Journal of Visualized Experiments. 2023;(196). doi:10.3791/64566

19. Arnold ND, Pickworth JA, West LE, et al. A therapeutic antibody targeting osteoprotegerin attenuates severe experimental pulmonary arterial hypertension. Nat Commun. 2019;10(1):5183. doi:10.1038/s41467-019-13139-9

20. Bray NL, Pimentel H, Melsted P, Pachter L. Near-optimal probabilistic RNA-seq quantification. Nat Biotechnol. 2016;34(5):525–527. doi:10.1038/nbt.3519

21. Melsted P, Ntranos V, Pachter L. The barcode, UMI, set format and BUStools. Bioinformatics. 2019;35(21):4472–4473. doi:10.1093/bioinformatics/btz279

22. Melsted P, Booeshaghi AS, Liu L, et al. Modular, efficient and constant-memory single-cell RNA-seq preprocessing. Nat Biotechnol. 2021;39(7):813–818. doi:10.1038/s41587-021-00870-2

23. Wolf FA, Hamey FK, Plass M, et al. PAGA: graph abstraction reconciles clustering with trajectory inference through a topology preserving map of single cells. Genome Biol. 2019;20(1):59. doi:10.1186/s13059-019-1663-x

24. Wolock SL, Lopez R, Klein AM. Scrublet: Computational Identification of Cell Doublets in Single-Cell Transcriptomic Data. Cell Syst. 2019;8(4):281–291.e9. doi:10.1016/j.cels.2018.11.005

25. Butler A, Hoffman P, Smibert P, Papalexi E, Satija R. Integrating single-cell transcriptomic data across different conditions, technologies, and species. Nat Biotechnol. 2018;36(5):411–420. doi:10.1038/nbt.4096

26. Lopez R, Regier J, Cole MB, Jordan MI, Yosef N. Deep generative modeling for single-cell transcriptomics. Nat Methods. 2018;15(12):1053–1058. doi:10.1038/s41592-018-0229-2

27. Traag VA, Waltman L, van Eck NJ. From Louvain to Leiden: guaranteeing well-connected communities. Sci Rep. 2019;9(1):5233. doi:10.1038/s41598-019-41695-z

28. Becht E, McInnes L, Healy J, et al. Dimensionality reduction for visualizing single-cell data using UMAP. Nat Biotechnol. 2019;37(1):38–44. doi:10.1038/nbt.4314

29. McInnes L, Healy J, Saul N, Großberger L. UMAP: Uniform Manifold Approximation and Projection. J Open Source Softw. 2018;3(29):861. doi:10.21105/joss.00861

30. Gillis-Buck E, Miller H, Sirota M, et al. Extrathymic Aire-expressing cells support maternal-fetal tolerance. Sci Immunol. 2021;6(61). doi:10.1126/sciimmunol.abf1968

31. Kudlova N, De Sanctis JB, Hajduch M. Cellular Senescence: Molecular Targets, Biomarkers, and Senolytic Drugs. Int J Mol Sci. 2022;23(8):4168. doi:10.3390/ijms23084168

32. Dodig S, Čepelak I, Pavić I. Hallmarks of senescence and aging. Biochem Med (Zagreb). 2019;29(3):483–497. doi:10.11613/BM.2019.030501

33. Ferrero R, Rainer P, Deplancke B. Toward a Consensus View of Mammalian Adipocyte Stem and Progenitor Cell Heterogeneity. Trends Cell Biol. 2020;30(12):937–950. doi:10.1016/j.tcb.2020.09.007

34. Christodoulides C, Lagathu C, Sethi JK, Vidal-Puig A. Adipogenesis and WNT signalling. Trends Endocrinol Metab. 2009;20(1):16–24. doi:10.1016/j.tem.2008.09.002

35. Hepler C, Shan B, Zhang Q, et al. Identification of functionally distinct fibro-inflammatory and adipogenic stromal subpopulations in visceral adipose tissue of adult mice. Elife. 2018;7. doi:10.7554/eLife.39636

36. Sun K, Tordjman J, Clément K, Scherer PE. Fibrosis and adipose tissue dysfunction. Cell Metab. 2013;18(4):470–477. doi:10.1016/j.cmet.2013.06.016

37. Tello D, Balsa E, Acosta-Iborra B, et al. Induction of the mitochondrial NDUFA4L2 protein by HIF-1α decreases oxygen consumption by inhibiting Complex I activity. Cell Metab. 2011;14(6):768–779. doi:10.1016/j.cmet.2011.10.008

38. Kruzynska-Frejtag A, Machnicki M, Rogers R, Markwald RR, Conway SJ. Periostin (an osteoblast-specific factor) is expressed within the embryonic mouse heart during valve formation. Mech Dev. 2001;103(1-2):183–188. doi:10.1016/s0925-4773(01)00356-2

39. Hamilton DW. Functional role of periostin in development and wound repair: implications for connective tissue disease. J Cell Commun Signal. 2008;2(1-2):9–17. doi:10.1007/s12079-008-0023-5

40. O’Reilly S, Tsou PS, Varga J. Senescence and tissue fibrosis: opportunities for therapeutic targeting. Trends Mol Med. 2024;30(12):1113–1125. doi:10.1016/j.molmed.2024.05.012

41. Schafer MJ, White TA, Iijima K, et al. Cellular senescence mediates fibrotic pulmonary disease. Nat Commun. 2017;8(1):14532. doi:10.1038/ncomms14532

42. Khan T, Muise ES, Iyengar P, et al. Metabolic dysregulation and adipose tissue fibrosis: role of collagen VI. Mol Cell Biol. 2009;29(6):1575–1591. doi:10.1128/MCB.01300-08

43. Wolff G, Taranko AE, Meln I, et al. Diet-dependent function of the extracellular matrix proteoglycan Lumican in obesity and glucose homeostasis. Mol Metab. 2019;19:97–106. doi:10.1016/j.molmet.2018.10.007

44. Penke LR, Speth JM, Huang SK, Fortier SM, Baas J, Peters-Golden M. KLF4 is a therapeutically tractable brake on fibroblast activation that promotes resolution of pulmonary fibrosis. JCI Insight. 2022;7(16). doi:10.1172/jci.insight.160688

45. Zhang Y, Gai X, Li Y, et al. Autocrine GDF10 Inhibits Hepatic Stellate Cell Activation via BMPR2/ALK3 Receptor to Prevent Liver Fibrosis. Adv Sci (Weinh). 2025;12(19):e2500616. doi:10.1002/advs.202500616

46. Lin L, Han Q, Xiong Y, et al. Krüpple-like-factor 4 Attenuates Lung Fibrosis via Inhibiting Epithelial-mesenchymal Transition. Sci Rep. 2017;7(1):15847. doi:10.1038/s41598-017-14602-7

47. Ni C, Chen Y, Xu Y, et al. Flavin Containing Monooxygenase 2 Prevents Cardiac Fibrosis via CYP2J3-SMURF2 Axis. Circ Res. Published online July 5, 2022:101161CIRCRESAHA122320538. doi:10.1161/CIRCRESAHA.122.320538

48. Weinstock A, Brown EJ, Garabedian ML, et al. Single-Cell RNA Sequencing of Visceral Adipose Tissue Leukocytes Reveals that Caloric Restriction Following Obesity Promotes the Accumulation of a Distinct Macrophage Population with Features of Phagocytic Cells. Immunometabolism. 2019;1. doi:10.20900/immunometab20190008

49. Nance SA, Muir L, Lumeng C. Adipose tissue macrophages: Regulators of adipose tissue immunometabolism during obesity. Mol Metab. 2022;66:101642. doi:10.1016/j.molmet.2022.101642

50. Misawa T, Wagner M, Koyasu S. ILC2s and Adipose Tissue Homeostasis: Progress to Date and the Road Ahead. Front Immunol. 2022;13. doi:10.3389/fimmu.2022.876029

51. Feuerer M, Herrero L, Cipolletta D, et al. Lean, but not obese, fat is enriched for a unique population of regulatory T cells that affect metabolic parameters. Nat Med. 2009;15(8):930–939. doi:10.1038/nm.2002

52. Biffi G, Tuveson DA. Diversity and Biology of Cancer-Associated Fibroblasts. Physiol Rev. 2021;101(1):147–176. doi:10.1152/physrev.00048.2019

53. Arpinati L, Scherz-Shouval R. From gatekeepers to providers: regulation of immune functions by cancer-associated fibroblasts. Trends Cancer. 2023;9(5):421–443. doi:10.1016/j.trecan.2023.01.007

54. Guccini I, Revandkar A, D’Ambrosio M, et al. Senescence Reprogramming by TIMP1 Deficiency Promotes Prostate Cancer Metastasis. Cancer Cell. 2021;39(1):68–82.e9. doi:10.1016/j.ccell.2020.10.012

55. Ghosh K, Capell BC. The Senescence-Associated Secretory Phenotype: Critical Effector in Skin Cancer and Aging. J Invest Dermatol. 2016;136(11):2133–2139. doi:10.1016/j.jid.2016.06.621

56. Wang Y, Apostolopoulou H, Sun IH, et al. Blocking Osteoprotegerin Reprograms Cancer Associated Fibroblast to Promotes Immune Infiltration into the Tumor Microenvironment. Preprint posted online October 17, 2025. doi:10.7554/eLife.107984.1

57. Suda M, Chaib S, Langhi Prata LGP, et al. Endothelial senescent-cell-specific clearance alleviates metabolic dysfunction in obese mice. Cell Metab. 2025;37(12):2455–2465.e6. doi:10.1016/j.cmet.2025.10.009

58. De Leon-Oliva D, Barrena-Blázquez S, Jiménez-Álvarez L, et al. The RANK-RANKL-OPG System: A Multifaceted Regulator of Homeostasis, Immunity, and Cancer. Medicina (Kaunas). 2023;59(10). doi:10.3390/medicina59101752

