## Supplemental Data for "Osteoprotegerin-Enabled Immune Evasion of Pathological Adipose Stromal Cells Drives Metabolic Dysfunction in Obesity"

*equal contribution

‡corresponding author

#### Document S1

Supplemental Figure Legends S1–S4.

Figure S1 (related to Figure 1)

Figure S2 (related to Figure 3)

Figure S3 (related to Figure 4)

Figure S4 (related to Figure 5)

***<u>Supplemental Figure Legends</u>***

**Figure S1 (related to Figure 1). Genes transcriptionally regulated in non-immune AT stromal cells by maintenance on a low-fat CD as compared to chronic consumption of a HFD.** Differential gene expression analysis of non-immune ($CD45^{-}$ $C12FDG^{+}$) AT stromal cells across both the CD and HFD conditions, with volcano plot specifically showing transcriptionally up- and downregulated genes in mice fed the CD.

**Figure S2 (related to Figure 3): Marker-based annotation of immune cell clusters and AT macrophage remodeling during HFD feeding** (A) Dotplot showing the top 5 genes marking each of 16 clusters in the AT immune dataset. Marker genes were defined as significantly upregulated (Bayes factor > 3 and expressed in > 20% of cells within the respective cluster. Expression data are scaled between 0 (dark blue) and 1 (dark red). Dot size represents the fraction of cells within each cluster expressing each gene. **(B)** Matrixplot showing relative expression of LAM marker genes across different clusters. Expression data were normalized with z-score transformation. Blue and red represent the low and high expression of a gene, respectively, relative to the median expression level. (C) Time-dependent shifts in $CD163^{+}$ vs. $CD9^{+}$ macrophage abundance in mouse eWAT over 16 weeks of HFD consumption.

**Figure S3 (related to Figure 4). Integration of AT stromal and cancer stromal datasets identifies a shared iCAF/sAPC cluster (A)** UMAP of stromal cells colored by cluster identity (left) and by dataset of origin (right). (B) Barplot showing the percentage of cells from each dataset in each cluster. Most clusters primarily contain cells either from the cancer stromal or AT stromal datasets, except clusters 1 and 5. Cluster 5 contains both iCAFs and sAPCs. (C) Split violin plots showing expression of shared iCAFs and sAPCs genes, obtained by differential expression analysis of iCAFs vs. rest of cells in the cancer dataset and sAPCs vs. rest of cells in the AT non-immune stromal dataset.

**Figure S4 (related to Figure 5). OPG secretion by stromal progenitor cells from CD-fed vs. HFD-fed mice.** ELISA data, showing that AT stromal progenitor cells isolated and cultured from mice with DIO secrete substantially more OPG into the medium than do similar cells from CD-fed control mice (A) eWAT, (B) sWAT. *$p<0.05$, ***$p<0.001$ unpaired two-tailed Welch's t-test.

### Figure S1

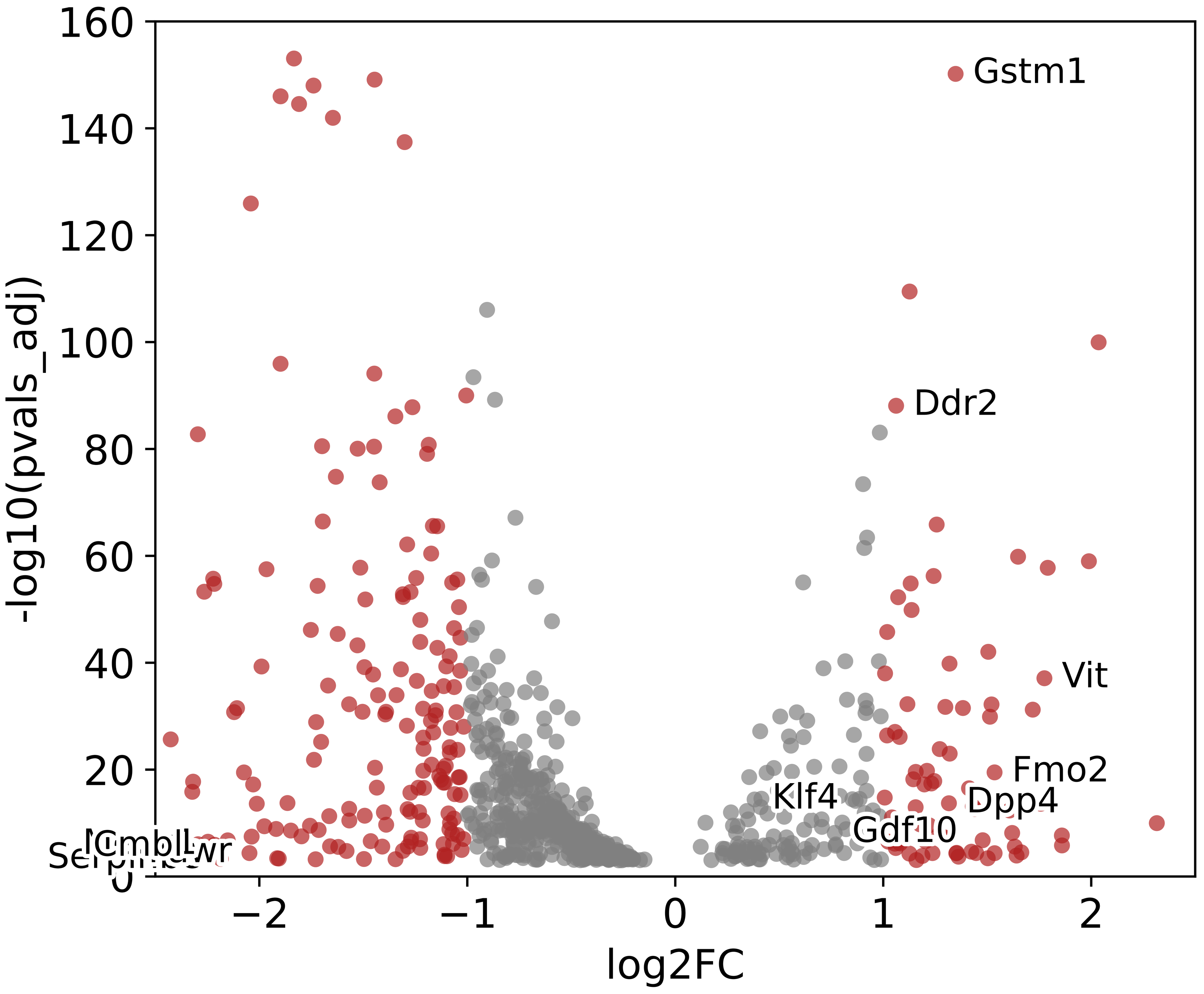
Gstm1
Ddr2
Vit
Fmo2
Dpp4
Klf4
Gdf10
160
140
120
100
80
60
40
20
0
−2
−1
0
1
2
-log10(pvals_adj)
log2FC

### Figure S2

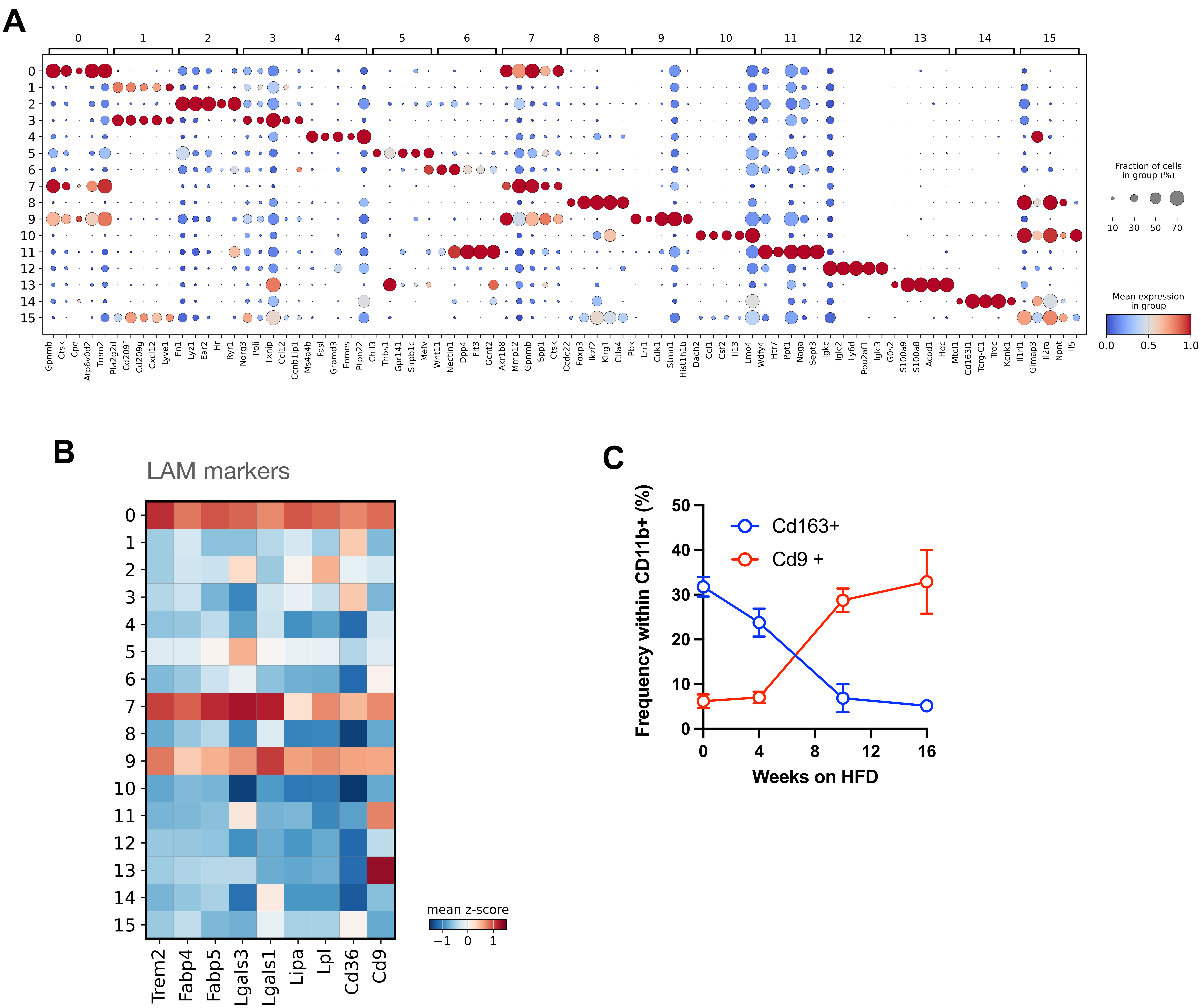

A
Fraction of cells in group (%)
10 30 50 70
Mean expression in group
0.0 0.5 1.0
Gpnmb Ctsk Cpe Atp6v0d2 Trem2 Pla2g2d Cd209f Cd209g Cxcl12 Lyve1 Fn1 Lyz1 Ear2 Hr Ryr1 Ndrg3 Poli Txnip Ccl12 Ccnb1ip1 Ms4a4b Fasl Gramd3 Eomes Ptpn22 Chil3 Thbs1 Gpr141 Sirpb1c Mefv Wnt11 Nectin1 Dpp4 Flt3 Gcnt2 Akr1b8 Mmp12 Gpnmb Spp1 Ctsk Ccdc22 Foxp3 Ikzf2 Klrg1 Ctla4 Pbk Lrr1 Cdk1 Stmn1 Hist1h1b Dach2 Ccl1 Csf2 Il13 Lmo4 Wdfy4 Htr7 Ppt1 Naga Sept3 Igkc Iglc2 Ly6d Pou2af1 Iglc3 G0s2 S100a9 S100a8 Acod1 Hdc Mctl1 Cd163l1 Tcrg-C1 Trdc Kcnk1 Il1rl1 Gimap3 Il2ra Npnt Il5
B
LAM markers
mean z-score
−1 0 1
Trem2 Fabp4 Fabp5 Lgals3 Lgals1 Lipa Lpl Cd36 Cd9
C
Cd163+
Cd9 +
Frequency within CD11b+ (%)
Weeks on HFD

### Figure S3

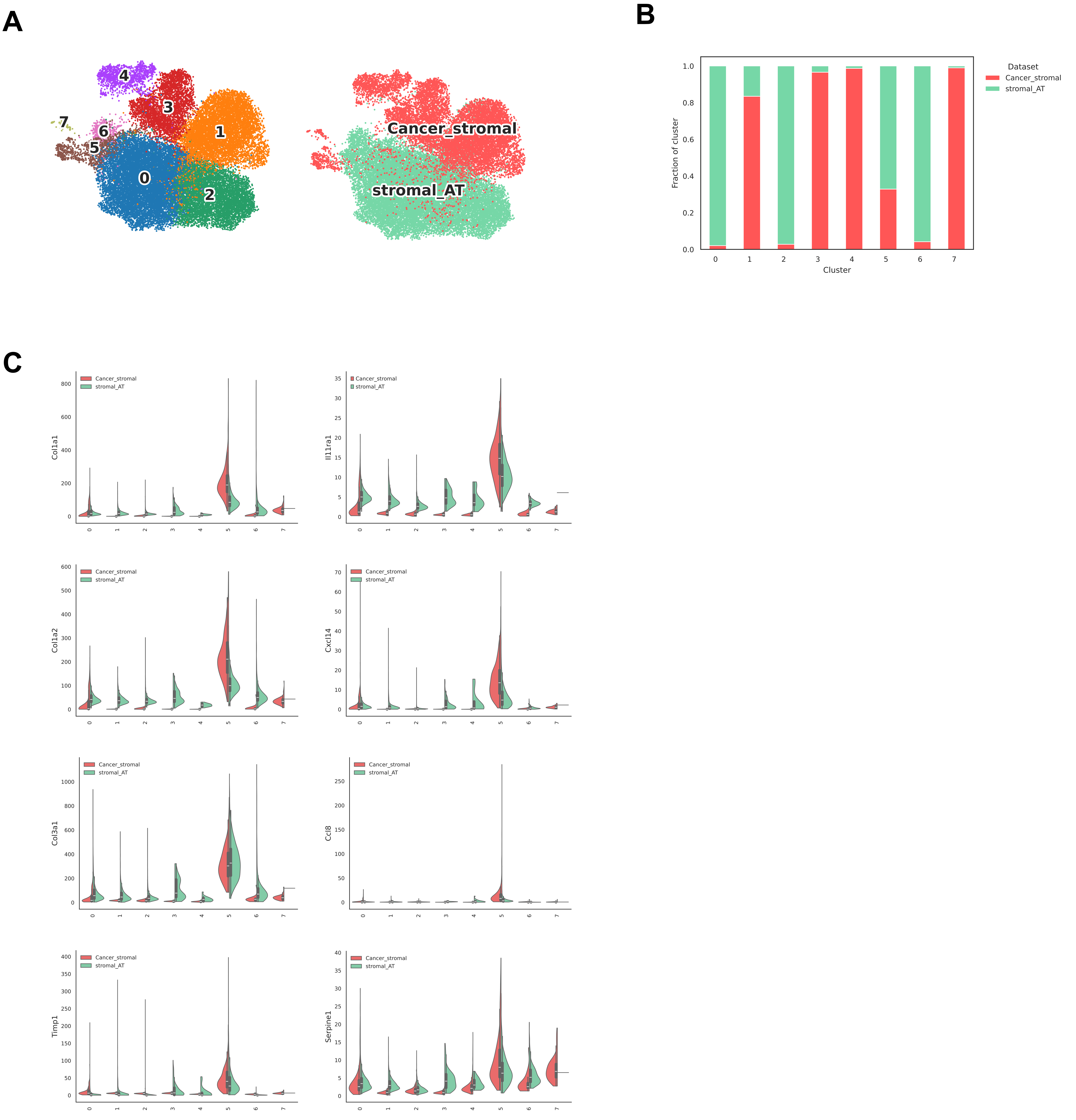
A
4
3
7
6
1
5
0
2
Cancer_stromal
stromal_AT
B
Dataset
Cancer_stromal
stromal_AT
Fraction of cluster
1.0
0.8
0.6
0.4
0.2
0.0
0 1 2 3 4 5 6 7
Cluster
C
Cancer_stromal
stromal_AT
Col1a1
Il11ra1
Col1a2
Cxcl14
Col3a1
Ccl8
Timp1
Serpine1

### Figure S4

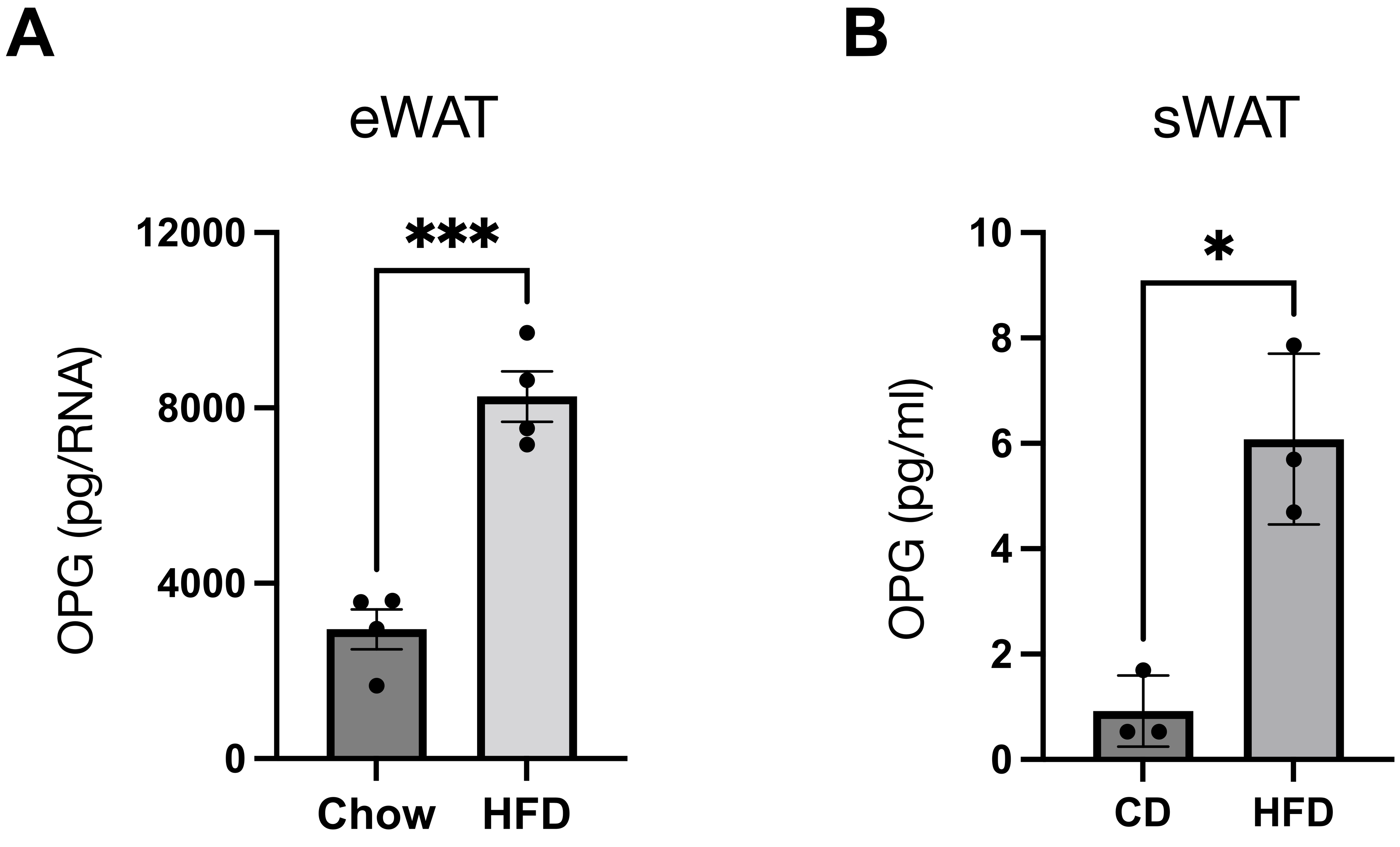
A
eWAT
OPG (pg/RNA)
12000
8000
4000
0
***
Chow
HFD
B
sWAT
OPG (pg/ml)
10
8
6
4
2
0
*
CD
HFD
